# Chronic Mild Stress Impairs Hippocampal Myelination through SOX6-Dependent Dysfunction of Oligodendrocyte Lineage Cells

**DOI:** 10.64898/2026.05.29.728722

**Authors:** Andrey Bombin, Cynthia Martin-Jimenez, Shun Yan, Shalini Saggu, Weiwei Liu, Yeimy Gonzalez-Giraldo, Emily Dew, Feng Zhou, Hanrui Wu, Quansheng Du, Tae Jin Lee, Ashok Sharma, Wei Zhang, Huidong Shi, Kai Jiao, Qin Wang

## Abstract

Chronic stress induces structural and functional changes in the brain, increasing susceptibility to major depressive disorder and other mental illnesses. Myelination deficits are a key pathological feature of stress-related disorders, yet the molecular mechanisms linking chronic stress to oligodendrocyte dysfunction remain poorly understood. Here, we used single-nucleus multiome sequencing to map gene expression and chromatin-accessibility remodeling in the hippocampus of mice exposed to chronic unpredictable mild stress (CUMS), a model that simulates key features of human daily stressors. CUMS induced broad molecular reprogramming across hippocampal cell populations, with stress-responsive gene networks significantly enriched for depression-associated genes. The oligodendrocyte lineage showed heightened vulnerability, with CUMS preferentially disrupting immature OPC/intermediate states and impairing OPC migration, OPC-to-ODC lineage progression, intercellular communication, and myelination. Integrated multiomic analysis identified stage-specific cis-regulatory elements and stress-sensitive gene regulatory networks, converging on SOXD transcription factors, particularly SOX5 and SOX6, as key regulators of OPC dysfunction. SOX6 ChIP-seq confirmed direct SOX6 binding at regulatory elements associated with genes controlling OPC morphogenesis, migration, and glutamatergic signaling. CUMS reduced SOX6 protein levels and SOX6-associated regulatory network activity in OPCs, whereas OPC-specific enhancer-driven restoration of SOX6 rescued stress-induced defects in OPC migration and myelination. Together, these findings define a stress-sensitive SOXD/SOX6 regulatory mechanism linking chronic stress to oligodendrocyte lineage dysfunction and myelination deficits, identifying SOX6 as a functional regulatory node with therapeutic potential for stress-related brain pathology.

## INTRODUCTION

Chronic stress induces widespread alterations in brain structure, function, and connectivity, increasing the risk for major depressive disorder (MDD) and other mental illnesses. The hippocampus, amygdala, and prefrontal cortex are particularly vulnerable to its effects. Chronic stress leads to reduced brain volume, diminished dendritic spines and synapses, impaired neural plasticity, and heightened neuroinflammatory responses in these regions, contributing to mood dysregulation and cognitive impairments ^1, 2, 3, 4, 5^. Despite growing recognition of stress-induced neurobiological damage, the underlying genetic and epigenetic mechanisms driving these alterations remain incompletely understood. This knowledge gap hampers the development of targeted therapeutic strategies, underscoring the need for a deeper mechanistic investigation into how chronic stress reshapes brain function at the molecular and cellular levels.

To better understand the mechanisms underlying morphological and functional alterations induced by chronic stress, bulk RNA sequencing and, more recently, single-nucleus RNA sequencing (snRNA-seq) have been used to profile transcriptomic changes in postmortem brain tissues from psychiatric patients, including those with MDD ^6, 7, 8^. These studies have revealed complex disease-associated transcriptional alterations. Notably, an snRNA-seq analysis of human postmortem prefrontal cortex identified excitatory neurons and oligodendrocyte lineage cells as particularly affected in MDD ^8^. While human postmortem studies provide valuable insights into the transcriptomic landscape of psychiatric disorders, challenges remain that limit the ability of these studies to elucidate the mechanisms driving these molecular changes. These challenges include limited sample availability, high inter-individual genetic and environmental variability, and the complexity of comorbid conditions. To address these limitations, animal models have served as powerful tools for dissecting the molecular drivers underlying stress-induced pathophysiology in controlled experimental settings. In mice, even brief exposure to acute stress can sufficiently alter transcriptomic features across different brain regions and cell types ^9, 10, 11^. A previous study using a 12-day unpredictable moderate stress model revealed gene signature changes associated with reduced excitatory neuronal activity in the prefrontal cortex ^12^, while studies on subchronic and chronic social defeat stress have identified elevated immune responses as a key feature of stress-related molecular changes ^13, 14, 15, 16^. Together, these studies provide valuable insights into the stress-induced molecular features across various brain cell populations and suggest transcriptomic profiles as sensitive signatures of different forms of stress. However, upstream regulatory mechanisms that drive these changes remain largely unresolved.

The rodent chronic unpredictable mild stress (CUMS) model simulates key features of daily stressors encountered in human life and induces behavioral and neurobiological changes resembling those observed in depressive disorders, including anhedonia, social withdrawal, cognitive impairment, reduced synaptic density and plasticity, deficits in neurogenesis, and glial dysfunction ^17, 18, 19^. Leveraging this model, we used a 6-week CUMS paradigm to determine how chronic mild stress reshapes transcriptomic and epigenomic landscapes in the hippocampus, a brain region highly susceptible to chronic stress and characterized by pronounced atrophic changes after prolonged exposure ^20, 21, 22, 23^. Despite this vulnerability, the cell-type-specific regulatory mechanisms linking chronic stress to hippocampal dysfunction remain poorly understood.

To address this gap, we performed single-nucleus multiome profiling, integrating RNA sequencing with Assay for Transposase-Accessible Chromatin sequencing (ATAC-seq) in the same nuclei, to resolve coordinated changes in gene expression and chromatin accessibility across hippocampal cell populations. By combining this approach with SOX6 chromatin immunoprecipitation sequencing (ChIP-seq) and functional validation, we mapped stress-sensitive gene regulatory programs and identified molecular drivers of oligodendrocyte lineage dysfunction. Our study highlights the oligodendrocyte lineage as a selectively vulnerable target of chronic stress and uncovers a SOXD-centered regulatory mechanism linking impaired OPC migration, disrupted lineage progression, altered intercellular communication, and reduced myelination. More broadly, these findings provide a mechanistic framework for understanding how chronic stress remodels hippocampal cell states and identify SOX6-associated regulatory programs as potential targets for mitigating stress-induced myelination deficits.

## RESULTS

### CUMS induces widespread transcriptional changes, with significant enrichment of depression-associated genes across hippocampal cell populations

Three-month-old C57BL/6 male mice underwent the CUMS procedure following a well-established protocol ^24^ with minor modifications (Fig. 1A, Supp. Table 1). While control mice gained weight over the 6-week period, most CUMS-exposed mice lost weight (Fig. 1B, C). CUMS also induced anhedonia-like behavior, as evidenced by a significant reduction in sucrose preference compared with controls (Fig. 1D). At the cellular level, CUMS significantly reduced synaptic density in the hippocampus (Fig. 1E–H). Together, these phenotypes are consistent with depression-related behavioral and neurobiological alterations observed in humans.

**Fig. 1.**
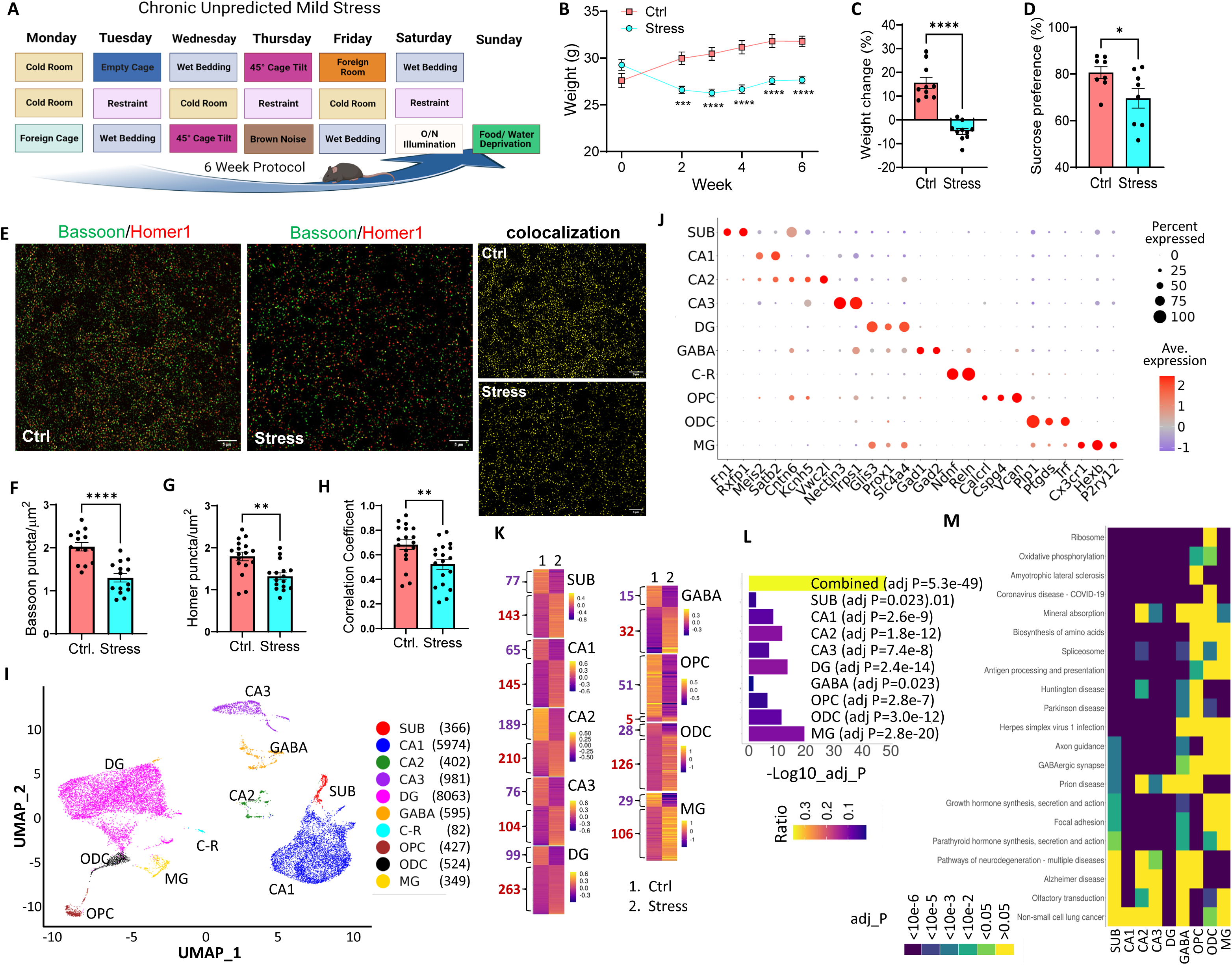
CUMS induces widespread transcriptomic alterations across hippocampal cell populations. **(A-D)** WT C57BL/6 male mice (3-month-old) underwent a 6-week chronic unpredictable mild stress (CUMS) protocol (A), with untreated littermates as controls. Body weight was measured weekly (B), and the percentage change after the 6-week protocol was calculated (C). At the end of the stress protocol, mice were assessed using the sucrose preference test (D). Data are presented as mean ± SEM. *, *p*<0.05; ***, *p*<0.001; ****, *p*<0.0001, by unpaired two-tailed Student’s *t* test. N=8-10 mice per group. **(E-H)** Hippocampal sections from control and stressed mice were subjected to immunofluorescence staining for Bassoon (presynaptic marker) and Homer1 (postsynaptic marker). Representative images of the CA1 stratum radiatum (SR) (E) and quantitation of Bassoon (F) and Homer1 (G) puncta, as well as their colocalization (H), are shown. Data are presented as mean ± SEM. **, *p*<0.01; ****, *p*<0.0001, by unpaired two-tailed Student’s *t* test. N =14-17 regions of interest (ROIs) from 4 mice per group. Scale bar, 5 µm. **(I-M)** Pooled hippocampal tissues from three stressed and three control mice were subjected to single-nucleus multiome analysis. **(I)** The UMAP chart showed nuclei clustered into multiple groups based on their gene expression profiles (snRNA-seq), with different brain cell types identified. The numbers in parentheses indicate the total number of nuclei (stress + control) in each cluster. **(J)** Dot plot showing the expression of known marker genes across cell types. **(K)** The heatmap summarizes the mean expression of significant DEGs between the control and stress groups in each cell population, with the number of altered genes indicated. Gene expression values in each group were scaled by subtracting the average expression per gene. **(L)** DEGs in each cluster, as well as across all clusters combined, are significantly enriched for genes associated with depression-related disorders based on the DisGeNET database. These include bipolar depression, depressive disorder, major depressive disorder, major depression (single episode), recurrent major depressive episodes, unipolar depression, and mental depression. **(M)** KEGG pathway activities were inferred for each cell using AUCell based on snRNA expression profiles. Differences in AUC scores between stress and control groups were evaluated using the Wilcoxon rank-sum test. The top three significantly enriched terms for each cell population are shown in the heatmap. SUB, subiculum neurons; CA1–3, pyramidal neurons in CA1–CA3; DG, dentate gyrus neurons; GABA, GABAergic neurons; C-R, Cajal-Retzius cells; OPC, oligodendrocyte precursor cells; ODC, oligodendrocytes; MG, microglia.

To investigate the molecular responses to CUMS in the hippocampus, a brain region highly susceptible to stress-induced damage ^20, 21, 22, 23^, we performed single-nucleus multiome analysis of chromatin accessibility and gene expression. Hippocampi from three control and three stressed mice were separately pooled and subjected to single-nucleus isolation followed by droplet-based snATAC-seq and snRNA-seq using the 10x Genomics platform. After quality control, 9,553 nuclei from control samples and 8,210 nuclei from stress samples were retained for downstream analyses. Unsupervised clustering of gene expression profiles identified at least 13 clusters (Supp. Fig. S1), which were assigned to 10 major cell types on the basis of known marker genes (Fig. 1I,J), including subiculum (SUB), CA1, CA2, CA3, dentate gyrus (DG), GABAergic neurons, Cajal-Retzius (C-R) cells, oligodendrocytes (ODCs), oligodendrocyte precursor cells (OPCs), and microglia (MGs). We identified significantly enriched marker genes for each cell type, containing known and novel cell type-specific markers (Supp. Table 2; Supp. Fig. S2). In addition, Area Under the Curve cell-level enrichment (AUCell) analysis ^25^ using the Kyoto Encyclopedia of Genes and Genomes (KEGG) database revealed signature biological pathways associated with individual cell populations (Supp. Fig. S3, Supp. Table 3).

We next identified differentially expressed genes (DEGs) between stress and control conditions for each cell population and found that CUMS induced broad transcriptomic alterations across all major cell types (Fig. 1K, Supp. Table 4). C-R cells were excluded from this analysis because of their low abundance (n = 82 combined nuclei). Given the strong link between chronic stress and depression, we compared the stress-induced DEG sets with depression-associated genes from the DisGeNET database. Notably, depression-associated genes were significantly enriched among the stress-responsive DEGs in every cell population, both individually and in the combined dataset (Fig. 1L). Gene Ontology (GO) term enrichment analysis further showed that many altered genes were associated with synaptic regulation, with enriched biological process, cellular component, and molecular function terms related to neurotransmitter receptors, transporters, and ion channels (Supp. Fig. S4, Supp. Table 5). These findings indicate that CUMS causes widespread transcriptomic dysregulation across hippocampal cell types, with prominent effects on synaptic gene programs and strong overlap with depression-associated genes.

To further determine how CUMS alters functional gene-set activity across individual cells or cell populations, we performed AUCell analysis using KEGG pathway gene sets (Fig. 1M, Supp. Fig. S5, Supp. Table 6). Consistent with the GO term analysis, multiple synaptic pathways, including glutamatergic, GABAergic, cholinergic, dopaminergic, and serotonergic synapse pathways, were significantly downregulated across neuronal populations (Supp. Fig. S5B). In addition, CUMS significantly altered gene sets associated with several neurodegenerative disorders, including spinocerebellar ataxia, prion disease, Parkinson disease, Huntington disease, and amyotrophic lateral sclerosis (Fig. 1M, Supp. Fig. S5A, Supp. Table 6). These findings suggest potential molecular connections between chronic stress and neurodegenerative processes and are consistent with clinical observations linking chronic stress to increased susceptibility to neurodegenerative diseases ^26, 27^.

### The myelination process is highly sensitive to CUMS and antidepressant treatment

We calculated the percentage of each cell population relative to the total nuclei in both control and stress groups. While the percentages of the majority of cell populations were comparable between the control and stress groups, the percentage of ODCs in the stress group was significantly lower than in the control group (Fig. 2A), suggesting that the oligodendrocyte lineage was particularly sensitive to CUMS. Consistently, we observed significantly reduced myelination in the hippocampus of the stress group compared to the control group, as indicated by a significantly reduced level of MBP immunoreactivity (Fig. 2B, C). This phenotype was accompanied by a higher number of PDGFRα+ OPCs in the stress group compared to the control group (Fig. 2D, E).

**Fig. 2.**
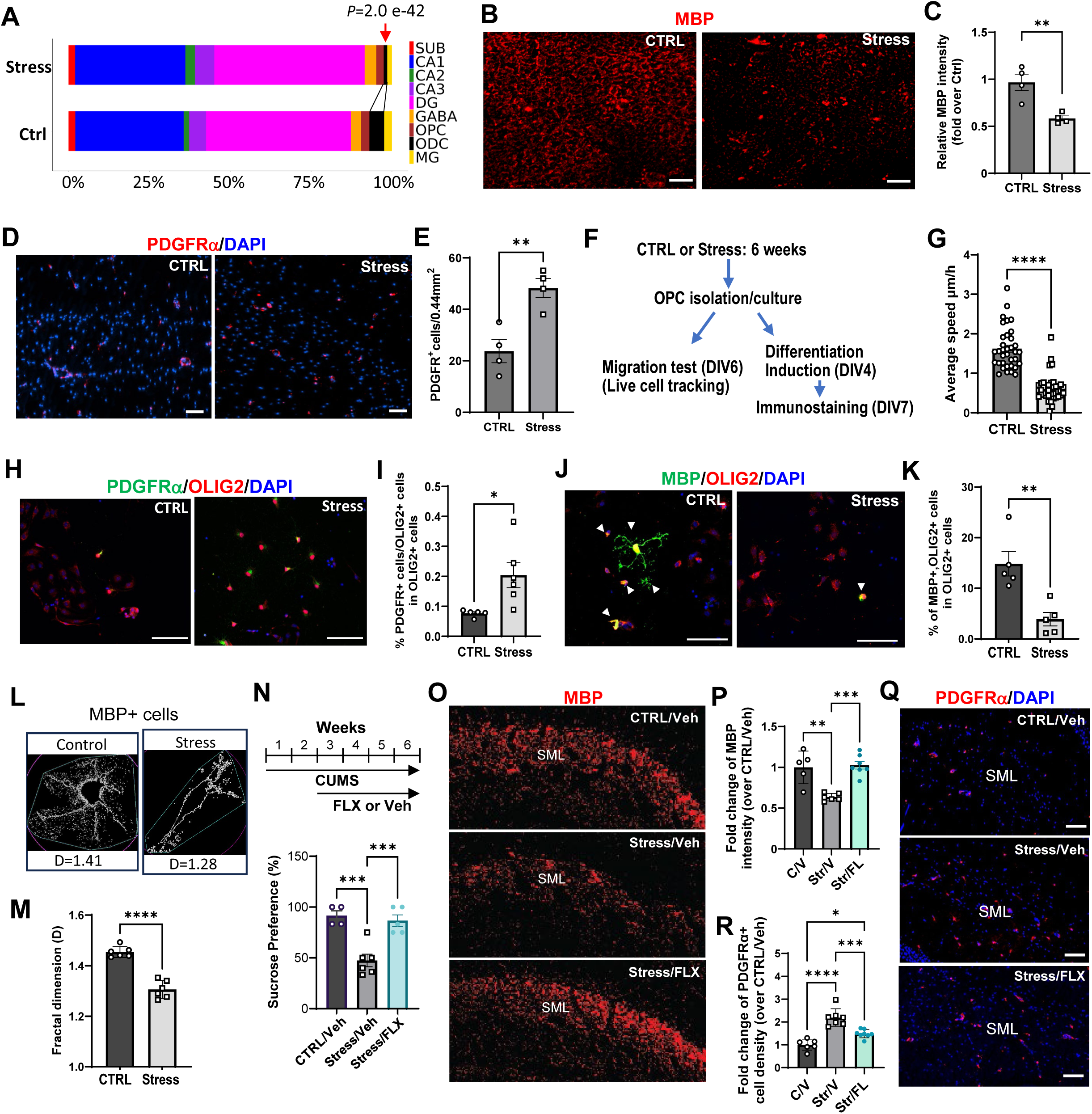
CUMS reduces myelination and impairs OPC migration and differentiation. **(A)** Bar graph showing the percentage distribution of nuclei across cell populations in the control and stress groups. The red arrow indicates the significant reduction in the ODC population in stressed mice relative to controls. *p*=2.0e-42, by one-sided two-proportion Z-test. **(B–M)** Brains were collected after 6 weeks of CUMS and processed for immunohistochemistry, or OPCs were isolated for culture-based assays. Age-matched littermates maintained under standard housing conditions served as controls. **(B,C)** Representative images (B) and quantification (C) of MBP immunoreactivity in the CA1 stratum lacunosum-moleculare (SLM). **(D,E)** Representative images (D) and quantification (E) of PDGFRα⁺ cells in the CA1 SLM. For (C,E), n=4 mice per group; *\*\*, p*<0.01 by unpaired two-tailed Student’s *t* test. **(F)** Schematic illustration of adult mouse-derived OPC culture, migration, and differentiation assays. OPCs isolated from control or stressed mice were plated in 24-well plates, and spontaneous migration was monitored on DIV6. In a separate set of cultures, differentiation was induced on DIV4, and cells were subjected to immunofluorescence staining on DIV7. **(G)** Quantification of average movement speed over a 24-hour period. N=35-40 cells from three independent cultures per group. *\*\*\*\*, p*<0.0001 by unpaired two-tailed Student’s *t* test. **(H,I)** Representative images (H) and quantification (I) of the percentage of PDGFRα⁺ cells. **(J,K)** Representative images (J) and quantification (K) of the percentage of MBP⁺ cells. **(L,M)** Representative images (L) and quantification (M) of fractal analysis performed on MBP⁺OLIG2⁺ cells to assess cell shape complexity. For (I,K,M), n=5-6 independent cultures per group; *, *p*<0.05; **, *p*<0.01; ****, *p*<0.0001 by unpaired two-tailed Student’s *t* test. **(N–R)** Mice subjected to CUMS received vehicle under regular housing conditions or fluoxetine (10 mg/kg/day, i.p.) under enriched housing conditions during weeks 3–6 of the CUMS procedure. **(N)** Sucrose preference test. **(O,P)** Representative images (O) and quantification (P) of MBP immunoreactivity in the CA1 SLM. **(Q,R)** Representative images (Q) and quantification (R) of PDGFRα⁺ cells in the CA1 SLM. For (N,P,R), n=4-6 mice per group; *, *p*<0.05; **, *p*<0.01; ***, *p*<0.001; ****, *p*<0.0001 by one-way ANOVA followed by Tukey’s multiple-comparisons test. All data are shown as mean ± SEM. Scale bars: 50 µm for B, D, O, and Q; 90 µm for H, J, and L.

To further investigate the OPC/ODC dynamics, we isolated and cultured PDGFRα+ OPCs from mice subjected to CUMS and from control mice without stress (Fig. 2F, Supp. Fig. S6A). Since OPC migration to target sites is a critical step for maturation and myelination ^28^, we first assessed spontaneous OPC migration in cultures. Despite being maintained under identical conditions, OPCs derived from stressed mice exhibited significantly reduced mobility compared to those from control mice (Fig. 2G, Supp. Fig. S6B,C). We then tested OPC differentiation *in vitro*. Following differentiation induction, a higher percentage of OLIG2+ cells remained PDGFRα+ in the stress group compared to the control (Fig. 2H,I, Supp. Fig. S6D). Concurrently, the stress group exhibited a significantly lower number of MBP+OLIG2+ cells compared to the control group (Fig. 2J,K, Supp. Fig. S6E). In addition, MBP+ ODCs from the stress group displayed reduced ramification complexity compared to those from the control group (Fig. 2L,M). These findings collectively provide strong experimental evidence that CUMS has detrimental effects on the differentiation processes of OPC/ODC lineage cells. Furthermore, our results show that the impact of stress persists even when cells from the CUMS group were cultured under the same conditions as control cells, supporting the notion that the effects of stress on the OPC/ODC lineage are mediated through persistent mechanisms, such as epigenetic alterations.

Given the high vulnerability of the myelination process to CUMS, we further tested whether it also responds to antidepressant treatment. Mice undergoing CUMS were treated with fluoxetine (i.p., 10 mg/kg/day) and housed in an enriched environment during weeks 3–6 of the CUMS procedure. This treatment rescued the anhedonia-like phenotype, as indicated by the sucrose preference test (Fig. 2N). In fluoxetine-treated mice exposed to CUMS, hippocampal MBP levels were significantly increased compared with vehicle-treated CUMS mice and were restored to levels comparable to those of unstressed controls (Fig. 2O,P). In addition, fluoxetine treatment significantly reduced the number of PDGFRα+ OPCs relative to vehicle treatment (Fig. 2Q,R). Collectively, these data suggest that myelination is highly sensitive to chronic stress and that a fluoxetine-based intervention combined with enriched housing can restore myelination, further supporting the clinical relevance of this CUMS mouse model.

### CUMS impairs OPC/ODC lineage dynamics and reduces differentiation and maturation

To better understand the effect of CUMS on OPC/ODC dynamics, we re-clustered the OPC and ODC populations using the Monocle 3 package ^29^ and performed trajectory/pseudotime analysis (Fig. 3A). We identified four subclusters along a single trajectory line (Fig. 3A). Based on the original Seurat clustering and marker gene expression (Fig. 3B, Supp. Table 7), subcluster 2 was defined as OPC-m3 (to distinguish from OPCs based on Seurat clustering), subcluster 1 as ODC-m3, and subclusters 3 and 4 as intermediate populations transitioning from the OPC to the ODC state (Fig. 3A,B). Population-specific marker analysis further showed that the intermediate population exhibited a distinct transcriptomic profile rather than simply representing a mixture of OPC and ODC gene expression patterns (Fig. 3C, Supp. Table 7).

**Fig. 3.**
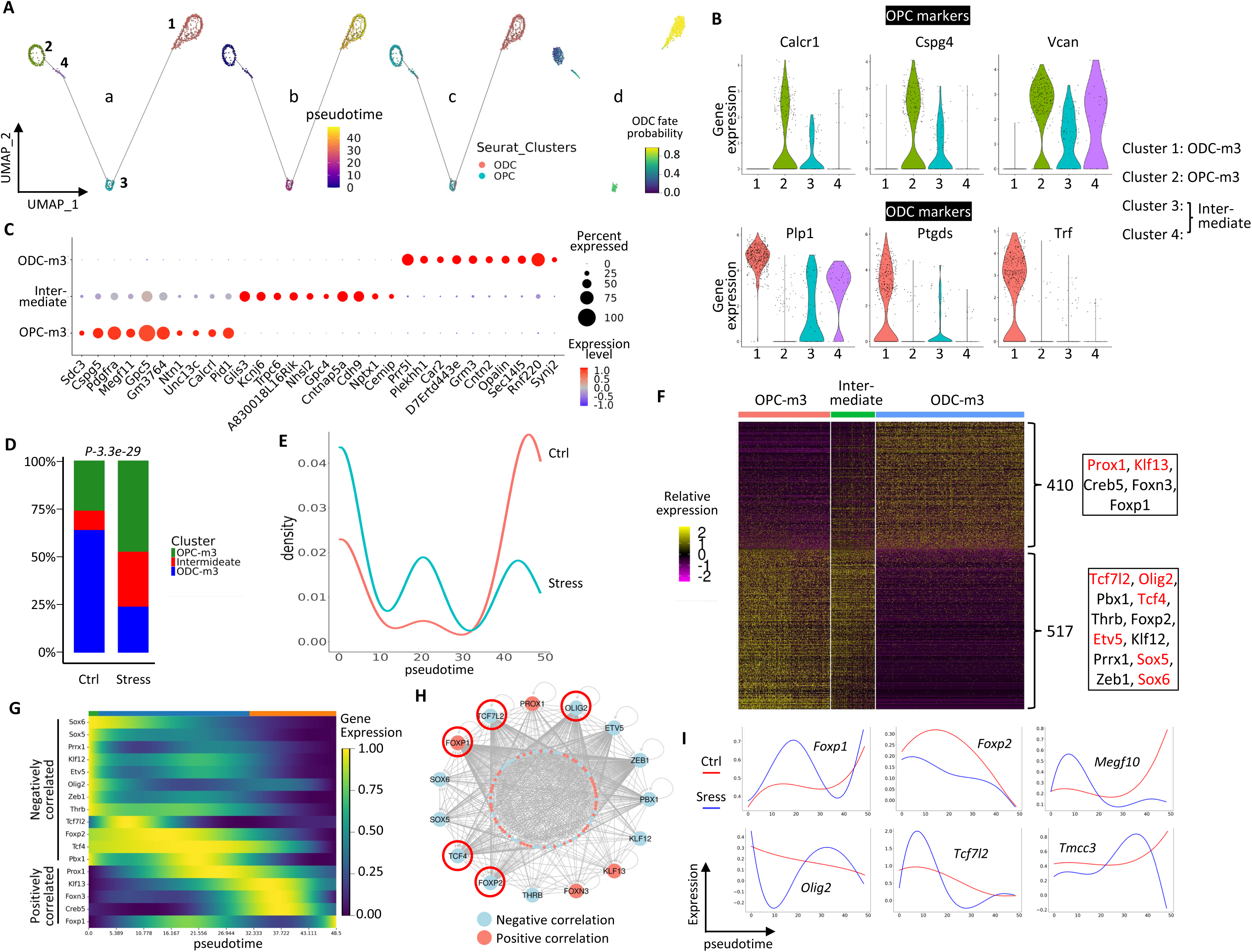
CUMS impairs OPC/ODC lineage dynamics. **(A)** Gene expression profiles of OPC and ODC nuclei, without separating control and stress conditions, were analyzed using Monocle 3 for trajectory inference. (a) UMAP plot of OPC and ODC nuclei showing four subclusters identified by Monocle 3. (b) The four subclusters are connected along a pseudotime trajectory, with cells colored according to pseudotime progression and the trajectory rooted at cluster 2. (c) The majority of nuclei in clusters 2, 3, and 4 were classified as OPCs by Seurat clustering, whereas all nuclei in cluster 1 were classified as ODCs. (d) Transition probability toward the ODC terminal state. **(B)** Violin plots showing the expression of known OPC and ODC marker genes across the four subclusters. Based on marker gene expression and pseudotime position, subcluster 1 was designated mature ODCs (ODC-m3), whereas cluster 2 was designated OPCs (OPC-m3). To distinguish these populations from the OPC and ODC clusters defined by Seurat analysis, they are labeled ODC-m3 and OPC-m3, respectively. Clusters 3 and 4 lie between OPC-m3 and ODC-m3 along the trajectory and express both OPC and ODC marker genes; these populations were therefore classified as intermediate cells. **(C)** Dot plot showing the top 10 marker genes for each cell population. **(D)** Bar chart showing the percentage of each cell population in control and stress groups. The distribution of cell populations differs significantly between the two groups (*p*=3.3e-29, Chi-square test). **(E)** Density plot showing the distribution of cells along the pseudotime trajectory in the control and stress groups. **(F)** Heatmap of 410 genes positively correlated and 517 genes negatively correlated with pseudotime progression. Among these, 5 TF genes were identified among the positively correlated genes and 12 among the negatively correlated genes. The TFs are listed in the box on the right; TFs shown in red are known regulators of OPC/ODC differentiation and function. **(G)** Expression patterns of the 17 TFs correlated with pseudotime during OPC-to-ODC differentiation. **(H)** Network analysis of pseudotime-correlated genes and their known upstream TFs. Known TF targets were obtained from the TFLink database and cross-referenced with pseudotime-correlated genes. The network plot shows 15 TFs (outer circle) and the top 50 positively and top 50 negatively correlated target genes (inner circle). CREB5 and PRRX1 were excluded because no target information was available in TFLink, likely due to limited publicly available ChIP data. TFs outlined in red represent those whose correlation with pseudotime was lost under CUMS. **(I)** Representative genes, including four TFs, showing loss of correlation with pseudotime under stress (blue) compared with control (red).

We next compared the subcluster compositions of OPC/ODC lineage cells between control and CUMS groups. In the stress group, the proportions of OPC-m3 and intermediate cells were increased, whereas the proportion of ODC-m3 cells decreased relative to controls (Fig. 3D). Consistent with this shift, pseudotime density analysis showed increased accumulation of precursor and intermediate populations and reduced representation of mature cells under stress (Fig. 3E). Together, these findings indicate that CUMS disrupts OPC/ODC lineage homeostasis and impairs progression from OPCs to mature ODCs, consistent with the marker-based analysis shown in Fig. 2.

To further define the transcriptional programs associated with OPC-to-ODC differentiation, we identified 410 genes positively correlated and 517 genes negatively correlated with pseudotime progression (*FDR < 0.05*; Fig. 3F, Supp. Fig. S7, Supp. Table 8A). Among these genes were 17 transcription factor (TF) genes, including known regulators of oligodendrocyte lineage development, such as *Sox5*, *Sox6*, and *Olig2* (Fig. 3F,G). To test whether these TFs act coordinately to drive the OPC-to-ODC differentiation program, we cross-referenced their known targets from TFLink ^30^ with the 927 pseudotime-correlated genes, restricting the analysis to targets supported by ChIP evidence. Notably, 864 of the 927 genes were linked to 15 of these TFs, and 844 were targeted by at least two TFs (Fig. 3H, Supp. Table 8B).

We then tested how CUMS affects this differentiation-associated transcriptional program. In the stress group, 244 pseudotime-correlated genes, including five of the 17 TFs, lost significant correlation with pseudotime (*FDR>0.1*, Supp. Table 8A). Representative trend plots, including those for TFs (*Foxp1*, *Foxp2*, *Olig2*, and *Tcf7l2*), illustrate stress-associated disruption of normal expression trajectories (Fig. 3I). Collectively, our data support the existence of a coordinated TF-driven transcriptional network underlying OPC-to-ODC differentiation, and this program is broadly disrupted by chronic stress, leading to accumulation of immature oligodendrocyte lineage cells in the hippocampus.

### CUMS disrupts intercellular communication between the OPC/ODC lineage and other cell types

We next analyzed differential gene expression in OPC/ODC lineage subpopulations between control and stress groups. Approximately 75% of DEGs were downregulated in OPC-m3 and intermediate cells, whereas more than 60% were upregulated in ODC-m3 cells, indicating differential effects of CUMS on immature versus mature oligodendrocyte lineage cells (Fig. 4A, Supp. Table 9). Consistent with the migration and differentiation defects observed under stress, multiple signaling pathways well established in these processes, including PKA, CXCR4, netrin, Rho, Reelin, and myelination signaling, were downregulated in OPC-m3 and intermediate cells, as revealed by Ingenuity Pathway Analysis (IPA) (Fig. 4B, Supp. Table 10). In contrast, only one pathway reached significance in ODC-m3 cells. These findings suggest that immature OPC/ODC lineage cells are more vulnerable to chronic stress.

**Fig. 4.**
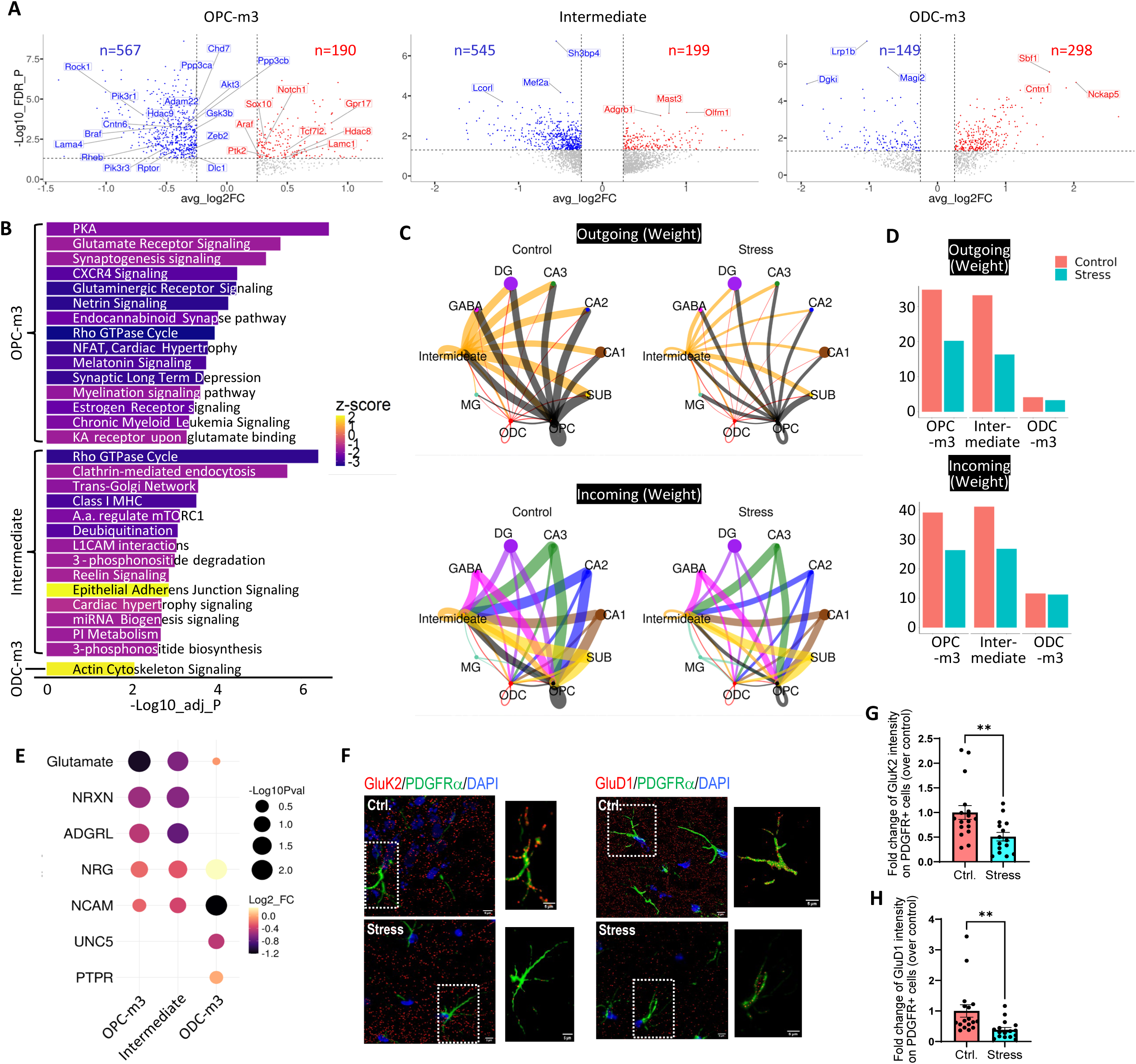
CUMS induces multiple molecular and cellular deficits in OPCs and ODCs. **(A)** Volcano plots showing DEGs between control and stress groups in OPC-m3, intermediate, and ODC-m3 cell populations. In OPC-m3 cells, genes involved in the myelination signaling pathway are labeled. In the intermediate and ODC-m3 populations, the three most significantly upregulated and downregulated genes are indicated. **(B)** DEGs were analyzed using Ingenuity Pathway Analysis (IPA) to identify significantly altered canonical pathways with a z-score > 1.5 or < -1.5 for each cell population. **(C)** Circle plots showing the weights of outgoing (top) and incoming (bottom) intercellular communication networks between OPC-m3, intermediate, or ODC-m3 cells and other cell populations under control (left) and stress (right) conditions. Node size reflects the number of cells in each population. **(D)** Bar graphs summarizing the total weights of outgoing (top) and incoming (bottom) intercellular communication associated with OPC-m3, intermediate, and ODC-m3 cells. **(E)** For each signaling pathway directed toward OPC-m3, intermediate, and ODC-m3 cells, information flow values were extracted from the CellChat analysis. Among the five pathways with the highest information flow in each cluster, the log2 fold change between stress and control conditions was calculated. Glutamatergic signaling showed the largest fold change in OPC-m3 cells and the second largest fold change in intermediate cells. **(F-H)** Hippocampal sections from control and stressed mice were subjected to immunofluorescence staining for GluK2, GluD1, and PDGFRα. **(F)** Representative images of the CA1 stratum lacunosum-moleculare (SLM). **(G,H)** Quantification of fold changes in GluK2 (G) and GluD1 (H) immunoreactivity in PDGFRα⁺ cells. N=15-17 sections from 4 mice per group. **, *p*<0.01 by unpaired two-tailed Student’s *t* test. Data are shown as mean ± SEM. Scale bar, 5 µm.

The OPC/ODC lineage cells actively communicate with other cell types to support normal neuronal and glial cell functions ^31^. We therefore analyzed how CUMS affects OPC/ODC intercellular communication networks using CellChat ^32^ (Supp. Fig. S8, S9). CUMS significantly reduced both incoming and outgoing signaling between OPC-m3 or intermediate cells and other cell populations, whereas changes in ODC-m3 cells were minimal (Fig. 4C, D, Supp. Fig. S10), consistent with the finding that fewer pathways were affected by CUMS in mature ODCs (Fig. 4B). We further examined the relative information flow into OPC-m3 and intermediate cells and identified multiple signaling pathways that were significantly altered by stress. Among the top five pathways, glutamatergic signaling, which is crucial for adaptive myelination ^33, 34^, showed the highest fold change in OPC-m3 and the second highest fold change in intermediate cells (Fig. 4E, Supp. Fig. S11), identifying it as a major target of stress in immature oligodendrocyte lineage cells. To further validate our bioinformatic finding, we examined the expression of two glutamate receptors known to regulate OPC differentiation and myelination ^35^, GluK2 (from the kainate receptor family) and GluD1 (from the delta receptor family), in PDGFRα+ OPCs in the hippocampus. The immunoreactivity of both receptors was significantly reduced in stressed mice compared with controls (Fig. 4F–H). Together, these data indicate that CUMS broadly disrupts intercellular communication involving OPCs, with glutamatergic signaling emerging as a prominent stress-sensitive pathway.

### Integrated snATAC-seq and snRNA-seq analyses reveal *cis*-regulatory elements that govern stage-specific gene expression in the OPC/ODC lineage

We next integrated the snRNA-seq and snATAC-seq datasets to identify *cis*-regulatory elements governing stage-specific gene expression in the OPC/ODC lineage. We first performed unsupervised clustering using the snATAC-seq data (Fig. 5A). Notably, cells assigned to the same identities based on snRNA-seq also clustered together on the snATAC-seq UMAP, supporting the notion that cells of the same type share not only similar transcriptomic profiles but also similar epigenetic states, as reflected by chromatin accessibility. We further performed weighted nearest neighbor (WNN) analysis ^36^ to generate an integrated UMAP from both snRNA-seq and snATAC-seq data (Fig. 5A). The relative positions of the clusters were largely preserved between the snRNA-seq UMAP and the WNN UMAP, further supporting the concordance between transcriptional and chromatin accessibility profiles and demonstrating the robustness of our single-nucleus multiome analysis across hippocampal cell populations.

**Fig. 5.**
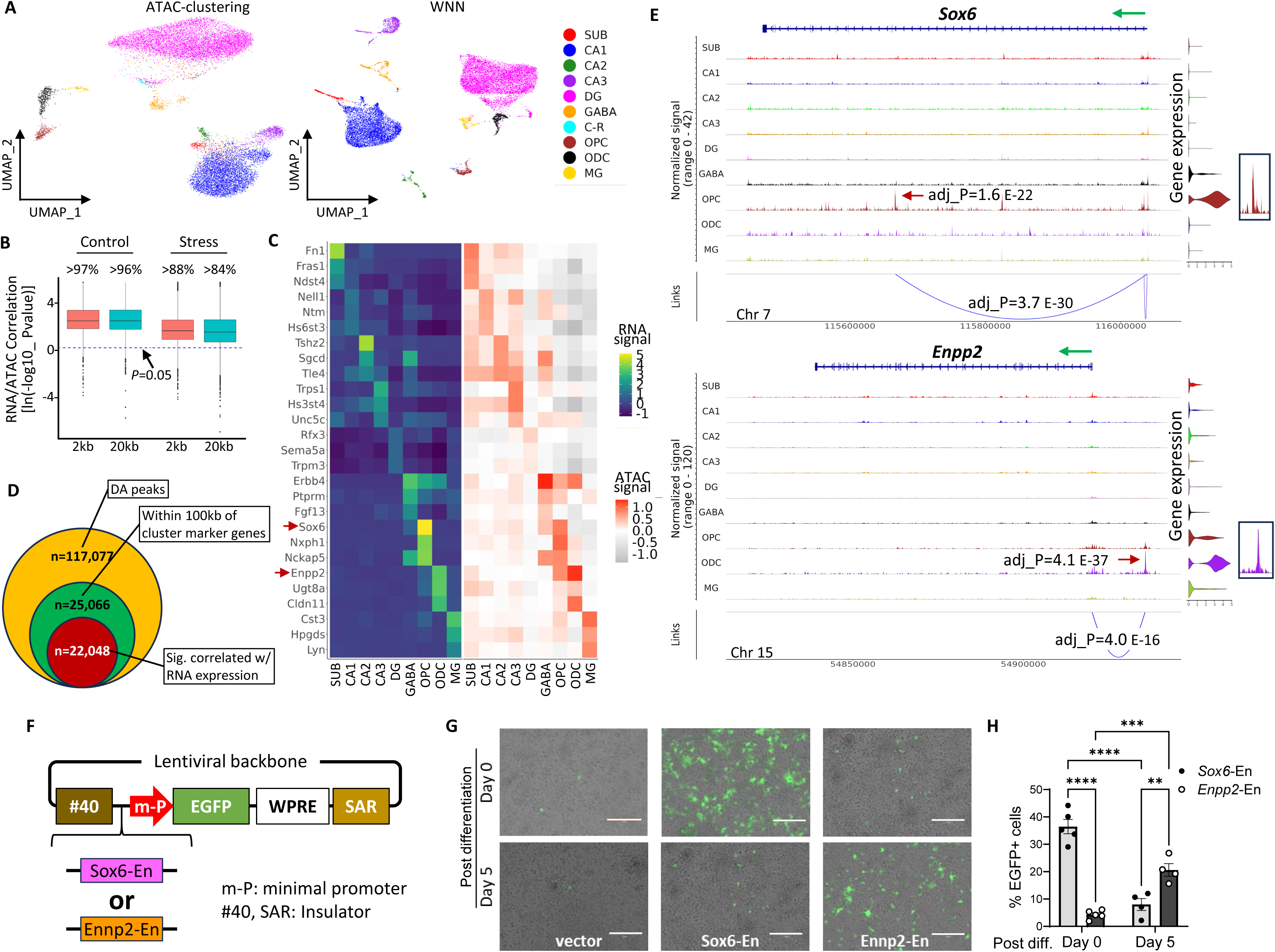
Identification of *cis*-regulatory elements governing cluster-specific gene expression by integrating snATAC-seq and snRNA-seq data. **(A)** The left UMAP plot shows clustering based on snATAC-seq data, and the right UMAP plot shows the results of weighted nearest neighbor (WNN) analysis integrating snATAC-seq and snRNA-seq data. In both plots, nuclei are colored according to the cell identities defined by the snRNA-seq analysis in Fig. 1I. **(B)** Correlations between chromatin accessibility and gene expression were calculated for all cluster marker genes. Chromatin accessibility was quantified using two genomic windows: from 2 kb upstream of the transcription start site (TSS) to the end of the gene body (“2 kb”), or across a 20 kb region flanking the gene body (“20 kb”). For visualization, the y-axis shows transformed P values as ln(-log10 *P* value). The dotted line indicates the significance threshold (*p*=0.05). Data are presented as median and interquartile range (IQR), with whiskers extending to 1.5xIQR. For most marker genes, chromatin accessibility was significantly correlated with gene expression. **(C)** Heatmap showing the mean expression of three top marker genes from each cluster whose chromatin accessibility was significantly correlated with their expression. Two color scales indicate the relative intensities of snRNA-seq and snATAC-seq signals. *Sox6* and *Enpp2* are highlighted by red arrows. **(D)** A total of 117,077 differentially accessible (DA) peaks were identified as showing significant differences in chromatin accessibility between a given cluster and all other cell populations (adj. *p*<0.05). Of these, 25,066 peaks were located within 100 kb of cluster-specific marker genes, and 22,048 showed significant correlations with expression of the associated marker genes (adj. *p*<0.05). **(E)** Coverage plots showing the significant DA peaks associated with *Sox6* (top panel, adj._*p*=1.6e-12) in OPCs and *Enpp2* (bottom panel, adj._*p*=4.1e-37) in ODCs. Accessibility of these peaks was also significantly correlated with *Sox6* and *Enpp2* expression, with adj. *p* values of 3.7e-30 and 4.0e-16, respectively. Green arrows indicate the transcription start site (TSS) and transcription direction of each gene. **(F)** Candidate enhancer regions from *Sox6* or *Enpp2* were inserted upstream of a minimal promoter in a lentiviral reporter construct. The #40 and SAR sequences served as insulators to minimize the influence of flanking genomic regions on GFP reporter expression. **(G, H)** OPCs were isolated from neonatal WT mouse brains and cultured in 24-well plates. On DIV2, cells were transduced with either an empty lentiviral reporter vector or reporter constructs containing the candidate *Sox6* or *Enpp2* enhancer elements. Forty-eight hours post-transduction, cells were induced to differentiate into ODCs. Images were acquired on days 0 and 5 after differentiation induction and are shown as merged bright-field and GFP fluorescence images. Representative images (G) and quantitation of the percentage of GFP+ cells (H) are shown. Scale bar, 50 µm. N=4-5 independent cultures per condition. Two-way ANOVA reveals significant effects for both differentiation day (*p*=0.0112) and enhancer type (*p*=0.0003), as well as their interaction (*p*<0.0001). **, *p*<0.01; ****, *p*<0.0001, by Tukey’s multiple comparisons test. Data are shown as mean ± SEM.

It is generally accepted that chromatin accessibility is concordant with gene expression; however, at least one recent study has reported that gene expression changes can occur independently of local chromatin accessibility changes ^37^. Therefore, we examined the relationship between these two features for cluster marker genes (Supp. Table 2). We calculated the correlation between gene expression and chromatin accessibility across two genomic regions: (1) from 2 kb upstream of the transcription start site (TSS) to the transcription termination site (TTS), and (2) from 20 kb upstream of the TSS to 20 kb downstream of the TTS. In both control and stress groups, the majority of marker genes showed significant correlations between expression and chromatin accessibility (Fig. 5B). Representative heatmaps of three marker genes from each cell type are shown in Fig. 5C. Intriguingly, the proportion of genes exhibiting significant concordance was reduced by approximately 10% in the stress group under both measurements (Fig. 5B), suggesting that CUMS partially disrupts the coupling between epigenetic regulation and gene expression.

Having established a strong correlation between chromatin accessibility and gene expression for cluster marker genes, we next asked whether integrated multiomic analysis could identify *cis*-regulatory elements associated with cluster-specific gene expression. We identified more than 117,000 differentially accessible (DA) peaks that showed significantly greater accessibility in one cluster than in all others (Supp. Table 11), with approximately 25,000 of these peaks located within 100 kb of cluster-specific marker genes (Supp. Table 12). Among these, approximately 22,000 peaks showed significant correlations with the expression of nearby cluster-specific marker genes, identifying them as candidate cluster-specific *cis*-regulatory elements (Fig. 5D, Supp. Table 13). We then compared these elements with known TF binding motifs in the JASPAR database (Supp. Table 14), with the top 10 enriched motifs for each cluster shown in Supp. Fig. S12A. This analysis identified motifs for TFs known to regulate lineage-specific development and function, including MEF2 factors in neurons ^38, 39^ and SOX10 and OLIG2 in the OPC/ODC lineage ^40, 41, 42^. We also performed de novo motif discovery using Hypergeometric Optimization of Motif EnRichment (HOMER), which identified additional enriched motifs that may represent novel regulatory elements involved in cell type-specific gene expression in the hippocampus (Supp. Fig. S12B).

To validate this approach experimentally, we focused on *Sox6* and *Enpp2*, whose expression is highly enriched in OPCs and ODCs, respectively ^42, 43, 44, 45^ (Supp. Fig. S2, Supp. Table 2). For each gene, we identified a significant DA peak in the corresponding cluster whose chromatin accessibility significantly correlated with gene expression (Fig. 5E). To determine whether these peaks function as enhancers driving stage-specific gene expression, we generated two lentiviral reporter constructs in which the candidate *Sox6* or *Enpp2* element was placed upstream of a minimal promoter. Two insulator sequences were included to minimize the influence of flanking genomic regions on EGFP reporter expression ^46^ (Fig. 5F). We then isolated OPCs from neonatal mice and transduced them with the *Sox6* reporter, the *Enpp2* reporter, or an empty vector control lacking an upstream enhancer. Before differentiation, the *Sox6* element, but not the *Enpp2* element, drove EGFP expression in OPCs. After differentiation, EGFP expression driven by the *Sox6* element was significantly reduced, whereas expression driven by the *Enpp2* element was significantly increased (Fig. 5G,H). These findings support the conclusion that the identified *Sox6* and *Enpp2* elements function as stage-specific enhancers and demonstrate that integrating snATAC-seq and snRNA-seq data can effectively identify *cis-*regulatory elements governing stage-specific gene expression in the OPC/ODC lineage. In addition, these elements provide useful tools for driving OPC- or ODC-specific gene expression, as demonstrated in the OPC-specific SOX6 rescue experiments below.

### SOXD regulatory network activity is downregulated by CUMS

To gain deeper insight into the transcriptional programs operating in distinct hippocampal cell populations and how they are altered by CUMS, we constructed gene regulatory networks (GRNs) using SCENIC+ (single-cell multiomic inference of enhancers and gene regulatory networks), which integrates snRNA-seq and snATAC-seq data with motif analysis to identify enhancer-driven regulons (eRegulons) ^47^. We identified eRegulons across all cell clusters (Supp. Table 15), with the five most abundant eRegulons for each cluster shown in Supp. Fig. S13. We then identified eRegulons whose activity was significantly enriched in individual clusters relative to the others (Supp. Table 16). For approximately 90% of the TFs associated with these eRegulons, their predicted binding motifs were also enriched in differentially accessible peaks located within 100 kb of a cluster marker gene (Supp. Tables 14 and 16), further supporting their roles in driving cluster-specific gene expression. Although neuronal populations shared broadly similar transcriptional networks, the strength and specificity of individual eRegulons varied among them, suggesting that distinct GRNs contribute to the transcriptional identities of different neuronal populations (Supp. Fig. S13). Notably, OPCs and ODCs displayed markedly distinct eRegulon patterns despite belonging to the same lineage, indicating substantial shifts in transcriptional regulation during lineage progression (Supp. Fig. S13).

In the next step, we compared eRegulon activity between control and stress conditions within each cell population and identified those significantly altered by CUMS (Fig. 6A, Supp. Fig. S14 and Supp. Table 17). CUMS affected both shared and cell type-specific regulatory programs: some stress-responsive eRegulons were altered across multiple cell populations (e.g. SETDB1-mediated eRegulon), whereas others showed selective changes in individual cell types. Supporting the disease relevance of these stress-responsive GRNs, nearly 40% of the 36 associated TFs are linked to mental disorders in DisGeNET (Fig. 6B), and genetic variations in *TCF4*, *SETDB1*, *BCL11A*, and *PHF21A*, have also been associated with altered gene expression and/or the onset of depression or schizophrenia in transcriptome-wide association studies ^48^ (Supp. Table 18).

**Fig. 6.**
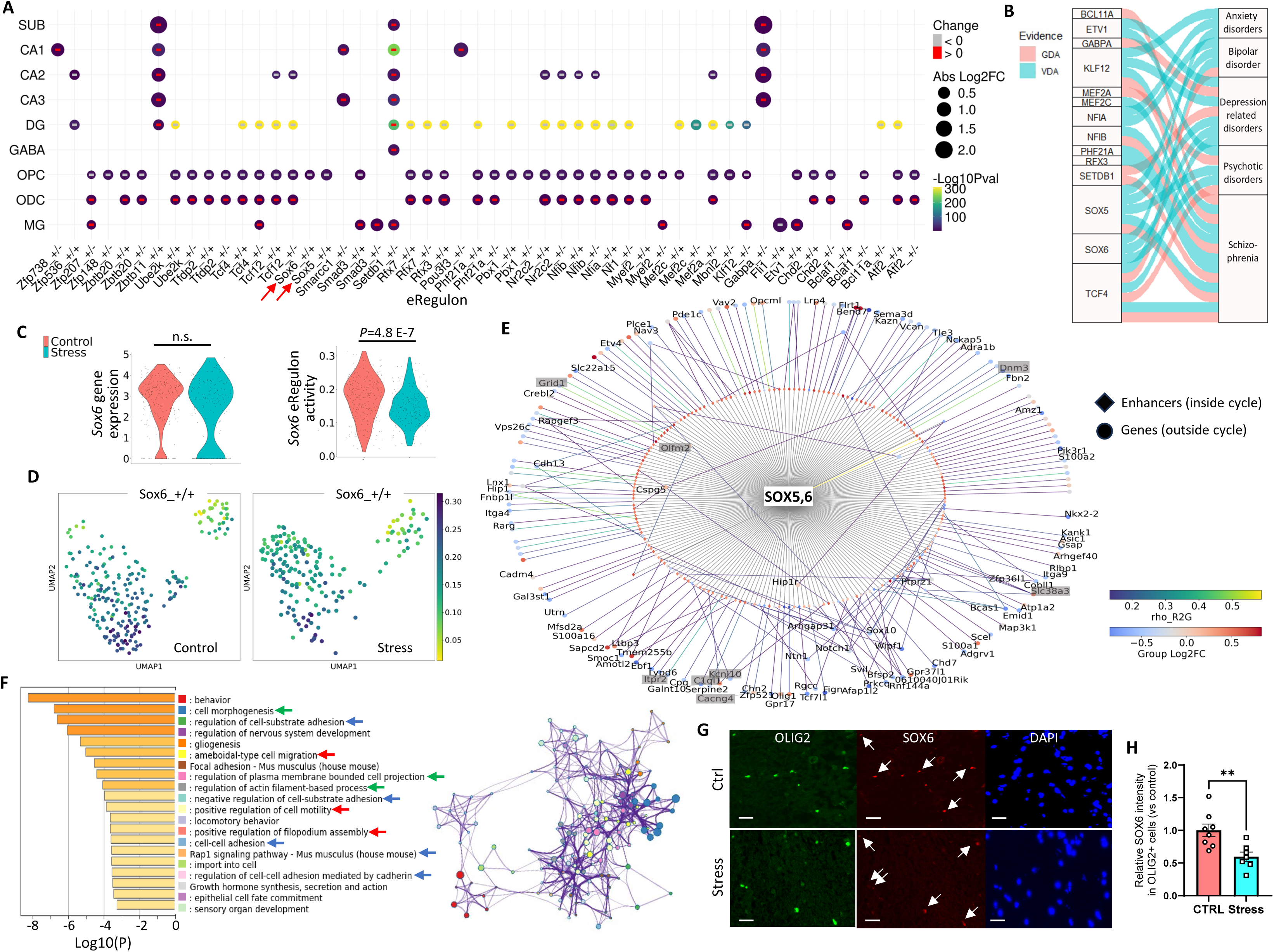
Reduced activity of SOXD-mediated eRegulons is associated with OPC defects induced by CUMS. **(A)** Dot plot summarizing eRegulons with significantly altered activity between control and stress groups within each cell cluster. **(B)** Fourteen of the 36 TFs driving the stress-responsive eRegulons shown in (A) are associated with mental disorders, based on cross-referencing with the DisGeNET database. **(C)** While *Sox6* transcript expression was not significantly altered in OPCs under stress (*left*), the activity of *Sox6*-mediated eRegulons was significantly reduced (*right*). **(D)** OPCs were re-clustered based on eRegulon activity. Feature maps illustrate decreased activity of *Sox6*-mediated eRegulons under stress at the single-nucleus level. **(E)** Network analysis revealed that *Sox5*- and *Sox6*-mediated eRegulons cooperate to regulate a complex gene network through their enhancers. The inner circle displays potential enhancers (snATAC peaks, represented by “♦”) linking *SOX5*/*SOX6* to their downstream targets, which are shown in the outer circle (represented by “●”). “Group Log2FC” quantifies the differential expression of target genes, and ‘rho_R2G’ represents the correlation coefficient between each candidate enhancer and its associated target genes. Genes that are OPC marker genes and/or significantly altered by stress are labeled. Eight genes involved in glutamatergic signaling and significantly downregulated in stressed OPCs are highlighted in gray boxes. **(F)** SOXD regulatory target genes identified by SCENIC+ were subjected to Metascape gene annotation and enrichment analysis. The left panel shows the top 20 significantly enriched terms, with terms related to cell migration, cell morphogenesis, and cell-cell/cell-matrix interactions highlighted by red, green, and blue arrows, respectively. The right panel shows the enrichment network of these terms. Each node represents an enriched term, with node size proportional to the number of associated genes. Terms with a similarity score > 0.3 are connected by edges, and edge thickness reflects the similarity score. **(G)** Hippocampal sections from control and stressed mice were subjected to immunofluorescence staining for OLIG2 (green) and SOX6 (red), with nuclei counterstained by DAPI (blue). Arrows indicate representative OLIG2⁺/SOX6⁺ double-positive cells. Scale bar, 20 µm. (**H**) Quantification of SOX6 immunoreactivity in OLIG2⁺/SOX6⁺ cells. N=7-8 sections from 4 mice per group, with at least 50 cells measured per mouse. **: *p*<0.01, by unpaired two-tailed Student’s *t* test. Data are mean ± SEM.

The SOXD family members, including SOX5 and SOX6, play a key role in regulating multiple processes in OPCs, including specification, migration, and terminal differentiation ^43, 49, 50^. The activities of eRegulons controlled by SOX5 (Supp. Fig. S15A) and SOX6 (Fig. 6C) were specifically reduced in OPCs under stress. This reduction is further illustrated by feature maps of the corresponding eRegulons (Fig. 6D, Supp. Fig. S15B). We reconstructed the regulatory network governed by these two TFs and found that they cooperatively regulate a complex gene network through enhancer-associated target genes (Fig. 6E, Supp. Table 19). Among the downstream targets of the SOX5/SOX6 eRegulons, at least eight genes, including *Grid1* (encoding GluD1), are implicated in glutamatergic signaling (highlighted in Fig. 6E). The CellChat analysis described above showed that incoming glutamatergic signaling to OPCs was significantly reduced under stress (Fig. 4E-H), suggesting that impaired SOX5/SOX6 eRegulon activity may contribute to this signaling deficit.

To further explore the functional relevance of this complex network, we performed MetaScape analysis ^51^ on the SOX5/SOX6-targeted genes to identify enriched biological pathways and GO terms (Fig. 6F). Among the top 20 significantly enriched terms, three were directly related to cell migration. Several additional terms were associated with cell morphogenesis and cell-cell or cell-matrix interactions, all of which are critical for cell migration (Fig. 6F). Together, these findings indicate that SOXD TFs regulate gene networks important for OPC identity, intercellular communication, and migration. To test whether CUMS reduces SOX6 eRegulon activity by regulating its expression, we examined *Sox6* at both RNA and protein levels. While *Sox6* transcript levels in OPCs were not altered by CUMS (Fig. 6C, left), SOX6 protein levels in OLIG2+ cells were significantly reduced in stressed mice relative to controls (Fig. 6G,H), suggesting that chronic stress regulates SOX6 through post-transcriptional mechanisms.

### SOX6 directly regulates genes controlling OPC morphogenesis and migration

To validate the potential targets of SOXD-family TFs identified by SCENIC+ (Fig. 6), we performed ChIP-seq in OPCs isolated from neonatal mice using a ChIP-grade SOX6 antibody ^52^. Peak calling with HOMER ^53^ identified 9,986 SOX6-associated peaks (Supp. Table 20). As shown in Fig. 7A,B, SOX6 binding was enriched primarily at promoter regions (±3 kb around TSSs). Substantial fractions of peaks were also located in intergenic and intronic regions (∼30% each), consistent with published SOX6 genomic distribution patterns ^50, 54^. Motif enrichment analysis revealed that SOXD-family motifs (SOX5/SOX6) were the top two enriched motifs within identified peaks (Fig. 7C). Motifs for additional SOX-family TFs were also significantly enriched, consistent with the conserved HMG DNA-binding domain and the shared 5′-ATTGTT core recognition sequence among SOX family members ^55, 56^. De novo motif discovery with HOMER further identified a highly significant motif closely matching the canonical SOX6 binding motif (Fig. 7D). These data provide strong support for the specificity of our SOX6 ChIP-seq assay.

**Fig. 7.**
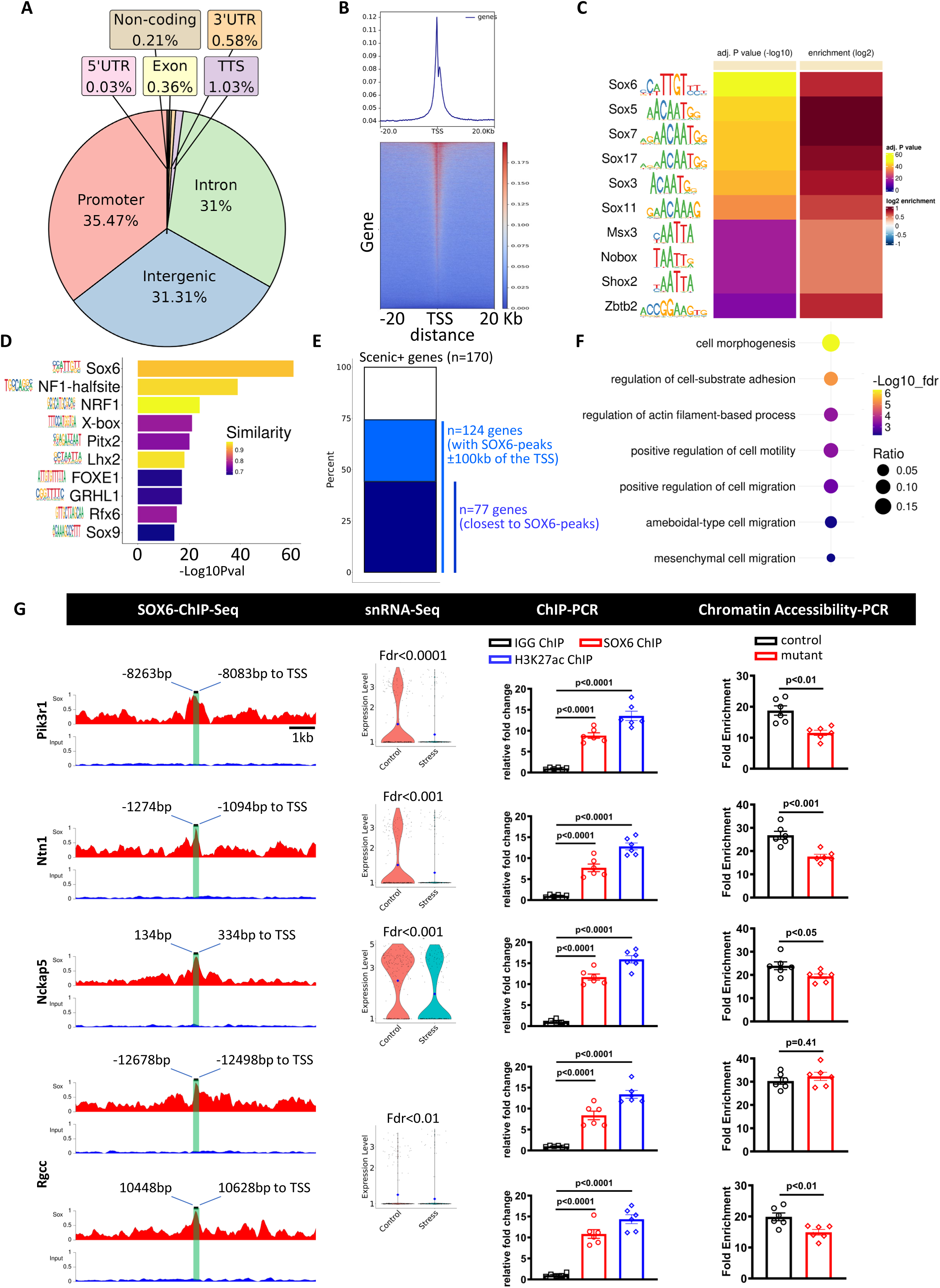
Identification of SOX6-bound regulatory elements associated with genes involved in cell morphogenesis and migration. **(A)** SOX6 ChIP-seq was performed using OPCs isolated from neonatal mice with two biological replicates. Peaks were called with HOMER, and the genomic distribution of SOX6 peaks is shown as a pie chart. **(B)** Distribution of SOX6 peaks around TSSs. **(C)** Top ten known motifs enriched within SOX6 peaks. **(D)** De novo motifs identified from SOX6 peaks; the top-ranked motif closely matches the canonical SOX6 motif. **(E)** Of the 170 genes identified by SCENIC+ as SOXD regulatory targets, 124 contain at least one SOX6 peak within ±100 kb of the TSS, and 77 of these 124 genes are the nearest genes to at least one SOX6 peak. **(F)** Enrichment analysis showing that terms related to cell morphogenesis and migration are significantly enriched among these 124 genes. **(G)** Summary of representative loci showing SOX6 ChIP-seq coverage tracks (average of two biological replicates), violin plots of differential gene expression in control and stress OPCs from snRNA-seq, ChIP-qPCR analysis of SOX6 and H3K27ac occupancy, and qPCR-based chromatin accessibility analysis in OPCs from control and stressed mice. For snRNA-seq analysis, the Wilcoxon rank-sum test was performed followed by Benjamini-Hochberg correction to control the false discovery rate (FDR). For ChIP-qPCR, one-way ANOVA followed by post hoc pairwise comparisons was used. For chromatin accessibility analysis, fold enrichment was calculated as the ratio of qPCR amplification from nuclease-treated DNA to that from the no-nuclease input control, and statistical significance was determined by unpaired two-tailed Student’s *t* test.

We next examined 170 genes predicted by SCENIC+ to be targets of the SOXD regulatory network (Fig. 6E). Of these, 124 contained at least one SOX6 peak within ±100 kb of the TSS, and 77 of these 124 genes were the nearest genes to at least one SOX6 peak (Fig. 7E, Supp. Table 21). Notably, all eight genes implicated in glutamatergic signaling that were highlighted in Fig. 6E, including *Grid1*, contained at least one SOX6 peak within ±100 kb of the TSS, supporting a direct role for SOX6 in regulating glutamatergic signaling in OPCs (Supp. Fig. S16).

Consistent with the pathway analysis shown in Fig. 6F, both GO terms and KEGG pathways related to cell morphogenesis and migration were significantly enriched among these 124 SOX6-peak containing genes (Fig. 7F). We selected five peaks associated with four genes for further validation (Fig. 7G) based on the following criteria: (1) the peaks were located within ±15 kb of the TSS; (2) the associated genes were downregulated under stress; (3) SOXD targeting was supported by both SCENIC+ prediction and SOX6 ChIP-seq; and (4) the genes are implicated in cell migration. ChIP-qPCR using primers specific to each peak confirmed SOX6 occupancy at these elements in isolated OPCs (Fig. 7G). Positive H3K27ac ChIP-qPCR signals further supported enhancer activity at these loci. Finally, chromatin accessibility analysis showed that four of the five elements exhibited significantly reduced accessibility in OPCs from stressed mice relative to controls, suggesting that stress induces epigenetic remodeling at SOX6-associated enhancers and thereby contributes to repression of their target genes. Together, our SOX6 ChIP-seq data, combined with SCENIC+ analysis of the multiomic datasets, indicate that SOXD-family factors directly regulate genes controlling OPC morphogenesis and migration, thereby contributing to OPC homeostasis and differentiation into ODCs.

### Downregulation of SOX6 underlies CUMS-induced defects in OPC migration and myelination

Given the important role of SOX6 in regulating OPC gene expression (Fig. 7) and its reduction by CUMS at the protein level (Fig. 6G,H), we next investigated whether decreased SOX6 contributes to the stress-induced defects in OPC migration and myelination. We first asked whether SOX6 levels are restored under conditions in which fluoxetine rescues the CUMS-induced myelination defects shown in Fig. 2N-R. In mice subjected to CUMS and treated with the fluoxetine-based intervention, SOX6 levels were significantly increased compared with those in vehicle-treated CUMS mice (Fig. 8A,B). These findings further validate that SOX6 levels are positively associated with myelination and support that SOX6 abundance is responsive to therapeutic intervention.

**Fig. 8.**
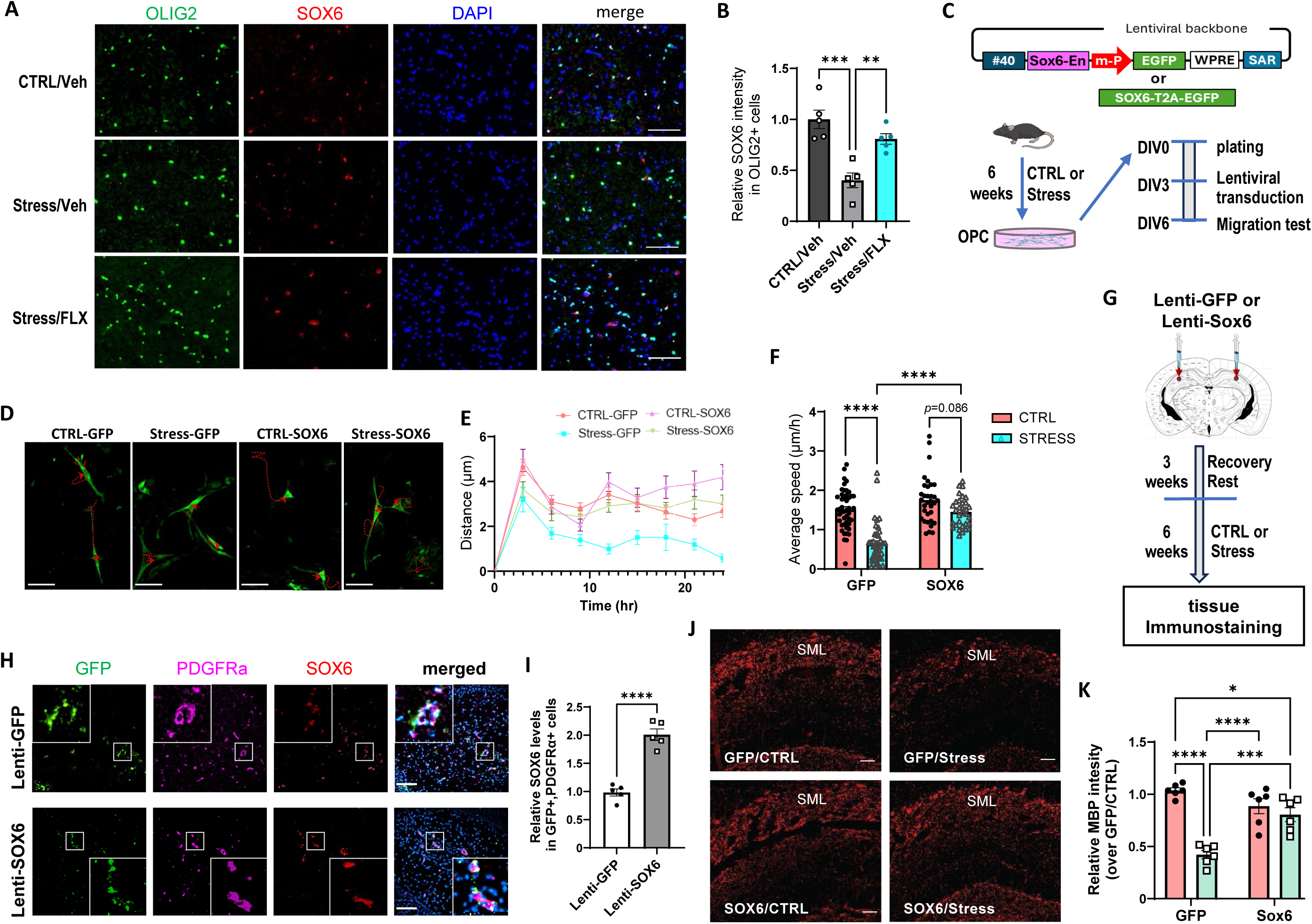
Elevating SOX6 levels in OPCs rescues stress-induced defects in OPC migration and myelination. **(A,B)** Fluoxetine treatment increased SOX6 levels. SOX6 immunofluorescence staining was performed on hippocampal sections from the indicated groups of mice treated as described in Fig. 2N. Representative images (A) and quantification (B) of SOX6 immunoreactivity in the CA1 SLM are shown. **, *p*<0.01; ***, *p*<0.001 by one-way ANOVA followed by Tukey’s multiple-comparisons test. **(C-F)** Exogenous SOX6 expression enhances migration of OPCs derived from CUMS mice. OPCs isolated from control and stressed mice were transduced with lentiviral constructs expressing GFP-T2A-SOX6 or GFP alone and then monitored for spontaneous migration (C). **(D)** Representative migration traces over 24 h. **(E,F)** Quantification of migration distance over 24 h (E) and average migration speed (F). N = 42–46 cells tracked from four independent cultures per group. Two-way ANOVA reveals significant effects for both stress treatment (*p*<0.0001) and SOX6 expression (*p*<0.0001), as well as their interaction (*p*=0.0054). ****, *p*<0.0001 by *post hoc* Fisher’s Least Significant Difference (LSD) multiple comparisons test. **(G)** Schematic of the *in vivo* rescue procedure. Lentiviral constructs expressing GFP-T2A-SOX6 or GFP alone were injected into the dorsal hippocampus. Three weeks after injection, mice were subjected to CUMS or left unstressed for 6 weeks. **(H,I)** Validation of lentiviral transduction of OPCs and SOX6 expression in the dorsal hippocampus CA1 SLM area at one-week post-injection. Hippocampal sections were immunostained for GFP, PDGFRα and SOX6. Representative images (H) and quantitation of SOX6 levels in GFP+PDGFRα+ cells (I) are shown. N=5 mice/group. ****, *p*<0.0001 by unpaired *t* test. **(J,K)** MBP immunofluorescence staining was performed on dorsal hippocampal sections from mice receiving lentiviral injection and subjected to control and CUMS, as indicated. Representative images of the CA1 SLM (J) and quantitation (K) of MBP immunoreactivity are shown. N=6 mice/group. Two-way ANOVA reveals significant effects for both stress treatment (*p*<0.0001) and SOX6 expression (*p*=0.0447), as well as their interaction (*p*<0.001). *, *p*<0.05; ***, *p*<0.001; ****, *p*<0.0001 by Tukey’s multiple comparisons test. All data are shown as mean ± SEM. Scale bars, 50 μm.

We next directly tested whether exogenous SOX6 expression in OPCs could rescue the migration and myelination defects induced by CUMS. To do so, we first introduced SOX6 into OPCs isolated from adult mice by lentiviral transduction using the OPC-specific *Sox6* enhancer identified in Fig. 5F-H (Fig. 8C). In OPCs derived from stressed mice, exogenous SOX6 expression significantly increased cell migration relative to GFP expression alone (Fig. 8D,E) and restored migration velocity to a level comparable to that of control cells (Fig. 8F). We further performed *in vivo* rescue experiments by introducing exogenous SOX6 into OPCs in the dorsal hippocampus of adult mice (Fig. 8G-I). In CUMS mice injected with Lenti-SOX6, myelination, as indicated by MBP staining, was significantly increased compared with that in mice injected with Lenti-GFP (Fig. 8J,K). Together, these *in vitro* and *in vivo* findings support that reduced SOX6 is a key mechanism underlying the CUMS-induced defects in OPC migration and myelination. More broadly, our findings identify SOXD family-regulated gene networks in OPCs as major targets of chronic stress.

## DISCUSSION

In this study, we used single-nucleus multiome sequencing, SOX6 ChIP-seq, and functional rescue experiments to define how chronic mild stress remodels the hippocampus at the transcriptomic, epigenomic, and cellular levels. We found that CUMS induces widespread molecular changes across hippocampal cell populations, with stress-responsive programs enriched for depression-associated genes. Among these populations, the OPC/ODC lineage showed particular vulnerability, including reduced myelination, impaired OPC migration and differentiation, disrupted intercellular communication, and altered progression along the OPC-to-ODC trajectory. Mechanistically, our analyses converged on a stress-sensitive SOXD regulatory program, centered on SOX5/SOX6 and especially SOX6, as a key contributor to OPC dysfunction. SOX6 ChIP-seq provided direct evidence that SOX6 binds regulatory elements associated with genes involved in OPC morphogenesis, migration, and glutamatergic signaling, strengthening the link between SOXD network disruption and stress-sensitive OPC phenotypes. Importantly, restoring SOX6 expression rescued CUMS-induced defects in OPC migration and myelination, moving SOX6 beyond a correlative regulatory signal to a functionally relevant mediator of stress-induced oligodendrocyte-lineage pathology. Together, these findings identify SOX6-dependent disruption of oligodendrocyte-lineage regulation as a major mechanism linking chronic stress to hippocampal myelination deficits.

Myelin damage is a key consequence of chronic stress and is commonly observed in major depression and other stress-related psychiatric disorders ^57^, yet the underlying molecular mechanisms remain poorly understood. In addition to displaying depression-related behavioral and neurobiological phenotypes, including reduced sucrose preference, body weight loss, and decreased hippocampal synaptic density ^17, 18, 19, 23, 58^, our CUMS model recapitulated reduced myelination, with OPC/ODC lineage cells showing heightened vulnerability to stress. Furthermore, the fluoxetine-based intervention, combined with enriched housing, rescued the myelination deficit and behavioral abnormalities in this model, highlighting the therapeutic relevance of our mechanistic investigation into chronic stress-induced myelin pathology.

Our data show that stress-induced defects are concentrated in immature oligodendrocyte lineage cells. OPC-m3 and intermediate cells displayed broad downregulation of pathways related to migration, differentiation, and myelination, whereas mature ODCs were less affected. Disruption of cell-cell communication was also confined to precursor cell populations (OPC-m3s and intermediate cells) but did not occur in mature ODCs. Intercommunication between OPCs and neurons, particularly through glutamate signaling, plays a crucial role in adaptive myelination, ensuring that myelin formation meets neuronal activity demands ^33, 34^. Our bioinformatic analyses identified glutamatergic signaling as a prominent target of chronic stress in OPCs, and this was supported by reduced expression of two glutamate receptors, GluD1 and GluK2 in hippocampal PDGFRα+ cells. These receptors are known to regulate OPC differentiation and myelination ^35^. In contrast to an earlier report ^15^ describing the presence of immune-oligodendrocytes in the prefrontal cortex of a repeated social defeat stress mouse model, we did not identify a similar immune-oligodendrocyte population in the hippocampus of our CUMS model. Our results suggest that chronic mild stress more strongly affects the synaptic regulatory function of the OPC/ODC lineage than immune activation. This discrepancy may also reflect differences in the brain regions examined, as our study focused on the hippocampus, whereas the previous study ^15^ investigated the prefrontal cortex.

Mechanistically, our multiomic analyses converged on SOXD-family regulatory networks, particularly SOX5 and SOX6, as stress-sensitive regulators in OPCs. SOX5 and SOX6 share the same DNA binding domain and are known to regulate multiple processes in oligodendrocyte progenitors, including specification, migration, and terminal differentiation ^43, 49, 50^. SOX5/SOX6 eRegulon activity was selectively reduced in stressed OPCs, and many predicted target genes were linked to cell migration, morphogenesis, and intercellular communication. SOX6 ChIP-seq then provided direct evidence that SOX6 binds regulatory elements linked to genes involved in OPC morphogenesis, migration, and glutamatergic signaling. Many SCENIC+-predicted SOXD targets were supported by nearby SOX6 peaks, and several validated SOX6-bound elements showed both enhancer-associated H3K27ac occupancy and reduced chromatin accessibility under stress. These findings move SOX6 beyond regulatory inference and support a model in which chronic stress disrupts OPC function in part by weakening SOX6-dependent control of genes required for migration, lineage progression, and intercellular communication.

Despite the importance of SOXD TFs in regulating OPC biology, it remains unclear whether their activity and expression can be altered by environmental stimuli and how such alterations contribute to stress-induced cellular dysfunction in OPCs. We observed a significant reduction in SOX6 protein levels in stressed mice compared to controls, despite unchanged RNA levels, suggesting that chronic stress targets SOX6 through post-transcriptional mechanisms. This distinction is important because it identifies SOX6 as a regulatory node whose activity can be selectively impaired without a corresponding change in transcript abundance. The functional importance of this pathway is underscored by the rescue experiments. Under the antidepressant treatment paradigm that restored myelination, SOX6 levels were also increased, suggesting that SOX6 is antidepressant-responsive and tracks with the myelination state *in vivo*. More directly, ectopic expression of SOX6 in OPCs derived from stressed mice restored migration, and *in vivo* SOX6 delivery to hippocampal OPCs enhanced MBP levels under CUMS. These findings move SOX6 from a correlative bioinformatic signal to a functionally relevant mediator of stress-induced OPC dysfunction. At the same time, the rescue is unlikely to imply that SOX6 acts alone. Rather, SOX6 appears to function as a pivotal node within a broader SOXD-centered regulatory architecture that integrates cell-intrinsic differentiation programs with cell-extrinsic signaling inputs. In this way, lower SOX6 levels may connect chronic stress to reduced glutamatergic signaling, impaired OPC migration, blocked differentiation, and, ultimately, decreased myelination.

It is also noteworthy that we used an OPC-specific *Sox6* enhancer to drive ectopic SOX6 expression. This stage-specific regulatory strategy was important for the *in vivo* rescue experiments because sustained SOX6 expression throughout the oligodendrocyte lineage prevents later maturation ^50^. Therefore, constitutive SOX6 expression driven by broadly active regulatory elements, such as the CMV enhancer/promoter, is not suitable for rescue experiments. In contrast, the endogenous *Sox6* enhancer identified in our study is active in OPCs but becomes attenuated after differentiation, enabling stage-restricted SOX6 expression. We identified this stage-specific enhancer by integrating gene expression and chromatin accessibility data and further validated it using reporter assays. Thus, this native *Sox6* enhancer provides a valuable tool for future OPC-specific studies.

Our results also have broader implications for psychiatric disease biology. When comparing the DEG list between stress and control groups with depression-related genes from DisGeNET, we found a significant enrichment of genes associated with depression-related disorders across all cell populations. Furthermore, nearly 40% of the TFs driving stress-induced changes in GRNs are linked to or implicated in various mental disorders. These findings reinforce the notion that chronic stress broadly reshapes the molecular landscape of the hippocampus in ways relevant to human psychiatric disease. At the same time, our study highlights the value of moving beyond neurons alone when considering the cellular basis of stress-related disorders. Although synaptic dysfunction remains a central feature of chronic stress, our data suggest that impaired oligodendrocyte lineage homeostasis, particularly at the precursor and transitional stages, is another major component of stress pathology. In addition to mental illness, our analysis of the DEGs between stress and control conditions also identified multiple terms related to neurodegenerative diseases. Clinical observations have shown that stress-associated disorders increase the risk of subsequent neurodegenerative diseases ^27^. The potential molecular connections between chronic stress and these neurological conditions identified here may help inform future studies on how chronic stress influences neurodegenerative disease risk or progression.

In summary, by combining single-nucleus multiome sequencing, SOX6 ChIP-seq, and functional validation in a CUMS model, our study reveals that chronic stress induces broad transcriptomic and chromatin-accessibility remodeling in the hippocampus and identifies the OPC/ODC lineage as a particularly vulnerable target. We show that chronic stress disrupts SOXD/SOX6 regulatory network activity, impairs OPC migration, intercellular communication, lineage progression, and myelination, and that restoring SOX6 rescues major stress-induced defects in OPC function and myelination. These findings provide a mechanistic framework for understanding how chronic stress alters oligodendrocyte-lineage biology and suggest that targeting SOXD-regulated programs may help mitigate stress-induced myelination deficits.

### Limitations of this study

First, we used the standard 10x Genomics protocol for single-nucleus isolation and did not recover clusters corresponding to astrocytes. This was not entirely unexpected, as astrocyte underrepresentation is a recognized limitation of snRNA-seq datasets generated using standard nuclei isolation protocols compared with single-cell approaches ^59, 60^. The absence of this cell type limited our ability to examine potential interactions between astrocytes and other hippocampal cell populations, including oligodendrocyte-lineage cells. Second, because of limitations in SOX6 antibody performance, the low yield of OPCs obtained from adult mice, and other technical challenges, we were unable to obtain reliable genome-wide ChIP-seq signals from adult OPCs. To overcome this limitation, we performed SOX6 ChIP-seq using OPCs isolated from neonatal mice. Therefore, it remains possible that SOX6 genomic occupancy differs between neonatal and adult OPCs. However, this concern is partially mitigated by SOX6 ChIP-qPCR validation in adult OPCs, which confirmed SOX6 occupancy at all five examined regulatory elements shown in Fig. 7G.

## Materials and Methods

### Mouse CUMS model and antidepressant treatment

WT C57BL/6 mice were used in the study. They were housed in the university animal housing facility, which provides a pathogen-free environment and is maintained on a 12-hour light/12-hour dark cycle. All animal experiments were conducted under a protocol approved by the Institutional Animal Care and Use Committees (IACUC) at Augusta University (Protocol number 2021-1061). All animal care and experimental procedures are reported in compliance with the ARRIVE guidelines ^61^.

The CUMS procedure was conducted following a well-established protocol ^24^ with modifications. Briefly, 3-month-old male mice were individually housed and exposed to a rotating sequence of stressors, which changed daily over a period of six weeks. The first week’s schedule is listed in Fig. 1A, and the full schedule is listed in Supp. Table 1. Control mice were group-housed under standard conditions. Body weight was recorded before the initiation of CUMS and then monitored weekly. At the end of the six-week procedure, mice underwent sucrose preference testing or were sacrificed for tissue collection.

For antidepressant treatment with fluoxetine, mice in the CUMS cohort were divided into two groups: one group received fluoxetine hydrochloride (catalog No. 0927; 10 mg/kg/day, i.p.), and the other received vehicle in 0.9% saline. Treatment began during the third week of the CUMS procedure. Concurrently, environmental enrichment (EE) was provided exclusively to the fluoxetine-treated group. EE consisted of larger cages containing additional corn cob bedding, nesting materials, and enrichment objects, including polycarbonate tunnels, balls, and housing domes. Treatments were administered daily from week 3 until the end of the experiment at week 6. At the end of the six-week procedure, mice underwent sucrose preference testing or were sacrificed for tissue collection.

### Sucrose preference test

Mice were assessed for sucrose preference using a two-bottle choice paradigm. Before testing, mice were habituated to two bottles containing water for 24 hours. They were then trained for 48 hours with access to one bottle containing water and one bottle containing sucrose solution. To minimize potential side preference, the positions of the water and sucrose bottles were switched after the first 24 hours of training. Following training, mice were tested for sucrose preference over a 24-hour period with free access to one bottle of water and one bottle of sucrose solution. During the test period, bottle positions were switched at 12 hours to control for side bias. Fluid intake from each bottle was measured, and sucrose preference (%) was calculated as [volume of sucrose solution consumed/(volume of water consumed + volume of sucrose consumed) × 100].

### SnRNA-seq analysis

Single nuclei were purified from the hippocampi of three control and three stressed male mice using the Chromium Nuclei Isolation Kit with RNase Inhibitor (PN-1000494, 10x Genomics). Libraries for snRNA-seq and snATAC-seq from the same nuclei were generated using the Chromium Next GEM Single Cell Multiome ATAC + Gene Expression Reagent Bundle and Chromium Next GEM Chip J Single Cell Kit (10x Genomics) on the Chromium X instrument (10x Genomics). Libraries were submitted to Azenta for deep sequencing, targeting at least 20,000 RNA-seq reads and 25,000 ATAC-seq reads per nucleus. Data analysis was performed using R v4.3.2 unless otherwise noted. Barcode-filtered files from the Cell Ranger ARC count pipeline were used to create Seurat v4.1.1 objects for the control and stress datasets. Nuclei expressing more than 400 unique molecular identifiers (UMIs), more than 200 features, and less than 5% mitochondrial UMIs were retained for further analysis. Potential doublets were identified with DoubletFinder v2.0.3 and removed from each dataset. Dataset integration and differential gene expression analysis were performed with Seurat v4.1.1. Each dataset was normalized using the “LogNormalize” method. Integration was based on canonical correlation analysis (CCA) anchors identified from the top 2,000 variable features. The integrated data were scaled and centered using a linear model in Seurat v4.1.1. Clustering analysis used 40 principal components, which were selected based on a clustering tree computed with Clustree v0.5.1, as well as JackStraw and elbow analyses in Seurat v4.1.1. Nuclei were clustered using a shared nearest-neighbor (SNN) graph and the Louvain algorithm, with an initial resolution of 0.2. Clusters were visualized using Uniform Manifold Approximation and Projection (UMAP) dimensionality reduction ^62^ in Seurat v4.1.1 and manually annotated based on marker gene expression visualized with feature plots. The UMAP visualization was generated with ggplot2 v3.4.4.

Genes showing positive differential expression in each cluster were identified using MAST, as implemented in Seurat v4.1.1 with MAST v1.28.0, and were restricted to genes expressed in at least 10% of nuclei in either group with a minimum log2 fold change of 0.25. Differentially expressed genes between stress and control groups were identified using the same method, without restricting the analysis to positive markers. P values were adjusted for multiple testing using the Bonferroni correction, as implemented in Seurat v4.1.1. Genes showing significant differential expression between stress and control groups, defined by adjusted P values < 0.05, were used for Kyoto Encyclopedia of Genes and Genomes (KEGG) pathway over-representation analysis, performed with clusterProfiler v4.10.0. In addition, clusterProfiler v4.10.0 was used to obtain gene sets for mouse KEGG pathways, which were then used to infer per-cell active gene sets with AUCell v1.24.0. Active gene sets present in at least 10% of nuclei in either group and showing significant differences between cell clusters or between stress and control groups within each cluster were identified using the Wilcoxon rank-sum test, as implemented in Seurat v4.1.1.

### Monocle 3 and CellRank analyses

For the analysis of OPCs and ODCs, cells from the corresponding clusters were extracted, and single-cell trajectory analysis was performed using Monocle3 v1.3.1. Briefly, the data were preprocessed by log normalization with size-factor adjustment, followed by dimensionality reduction to 100 components using principal component analysis (PCA). The percentage of variance explained by each component was plotted, and 85 dimensions were selected for downstream analysis. Control and stress datasets were aligned using the mutual nearest-neighbor algorithm to correct for batch effects. The data were then projected into a lower-dimensional space using UMAP, and new cell clusters were identified using the Leiden algorithm. Pseudotime was computed based on the principal graph and ordered to start from the OPC cluster. Relationships between pseudotime and gene expression were evaluated using a negative binomial model, as implemented in Monocle3 v1.3.1.

CellRank v2.0.6 was used to identify terminal states, their associated cells, and cell fate probabilities toward the ODC terminal state ^63^. Monocle3-derived UMAP coordinates from the OPC/ODC cell analysis described above were extracted and added to the Seurat object as a dimensional reduction, and Monocle3 pseudotime values were transferred to the Seurat object metadata. The updated Seurat object was then converted to AnnData (.h5ad) format using the as.anndata function implemented in scCustomize v3.2.4, retaining the normalized RNA matrix as the main layer, raw counts as an additional layer, and transferred dimensional reduction embeddings for downstream analysis.

A directed cell-cell transition matrix was then computed in CellRank using the pseudotime kernel, integrating transcriptomic neighborhood structure with Monocle3 pseudotime values to model directional state transitions ^63, 64^. Transition dynamics were visualized using the CellRank plot_projection function ^63^. The transition matrix was next used to initialize the Generalized Perron Cluster Cluster Analysis (GPCCA) estimator implemented in CellRank, and four macrostates were identified using the compute_macrostates function, with Monocle3 cell-cluster labels provided as guidance ^63^. Macrostates corresponding to OPC and ODC populations were designated as terminal states, and lineage fate probabilities toward these states were calculated using the compute_fate_probabilities function. Terminal macrostates and inferred fate probabilities were visualized using the plot_macrostates and plot_fate_probabilities functions, respectively. Lineage driver genes associated with the ODC terminal state were identified separately for control and stress groups using the compute_lineage_drivers function. Expression dynamics of the top 15 genes showing the strongest positive and negative correlations with ODC fate probabilities, and whose expression was also significantly associated with Monocle3 pseudotime as described above, were displayed using the CellRank heatmap function. Trajectory-specific expression trends for selected genes were modeled separately for control and stress cells using CellRank GAMR on Monocle3 pseudotime and visualized with the gene_trends function.

### CellChat analysis

Cell-cell interactions were inferred with CellChat v2.1.2 according to the package instructions ^65^. Briefly, for each group, normalized snRNA-seq count matrices were used to create a CellChat object. The default mouse ligand-receptor interaction database was used to infer cellular communication networks, with communication probabilities calculated using the computeCommunProb function and the “triMean” method. Communications involving cell groups with fewer than 10 cells were filtered out using the filterCommunication function. Pathway-level cell-cell signaling probabilities and aggregated communication networks were computed using the computeCommunProbPathway and aggregateNet functions, respectively, with default parameters. Preprocessed CellChat objects were merged using the mergeCellChat function. The aggregated cell-cell communication networks for incoming and outgoing signals were visualized using the netVisual_circle function. Edge width represents network weight, whereas vertex size represents the number of cells in each group. In addition, the summed incoming and outgoing interaction weights for OPC, intermediate, and ODC clusters were visualized as bar plots using ggplot2 v3.4.4. Differences in information flow between stress and control groups for each signaling pathway were extracted from the merged CellChat object using the rankNet function. For each cell cluster, signaling pathways were ranked based on the summed contribution scores from stress and control groups. The top five pathways for each cluster were visualized using ggplot2 v3.4.4.

### SnATAC-seq analysis and integration with snRNA-seq data

For the control and stress datasets, chromatin assay objects were generated from filtered feature matrices using Signac v1.12.0, and Seurat objects were created with Seurat v4.1.1. New peak sets were called using Signac v1.12.0 and MACS2 v2.2.6. For integration of the control and stress datasets, a common set of overlapping peaks was identified. Peaks longer than 10,000 bp or shorter than 20 bp were removed. The datasets were normalized using the term frequency-inverse document frequency (TF-IDF) method and integrated based on anchors identified through reciprocal latent semantic indexing, using all common peaks. In addition, for nuclei belonging to the OPC cluster that passed all quality-control filters, an OPC-specific peak set was called and used for differential peak analysis. Correlations between ATAC fragment counts within peaks of interest and gene expression, as well as fragment counts within the TSS region from −2.5 kb to +0.5 kb relative to the TSS, were evaluated using Spearman rank correlation tests, as implemented in base R. Differences in fragment counts within selected peaks between stress and control groups were assessed using the Wilcoxon rank-sum test, also implemented in base R. Genomic regions were visualized with genome-browser-style track plots using Signac v1.12.0. A UMAP integrating RNA-seq and ATAC-seq modalities was constructed from the weighted nearest-neighbor (WNN) graph, as implemented in Seurat v5.0.1, and visualized with ggplot2 v3.4.4.

### SCENIC+ analysis

Enhancer-driven gene regulatory networks were identified using the SCENIC+ pipeline v1.0a1 ^47^. Input data were prepared according to the SCENIC+ instructions. Briefly, snRNA-seq data were normalized by total expression and log-transformed using Scanpy v1.8.2. Highly variable genes were identified, the data were reduced to 50 dimensions by principal component analysis, and a nearest-neighbor graph was constructed with Scanpy v1.8.2 ^47, 66^. For each cell type, snATAC-seq peaks were called with Seurat v4.1.1 and MACS2 v2.2.6, followed by filtering to remove nonstandard chromosomes and mm10 blacklist regions (34, 63, 64). Region sets and a latent Dirichlet allocation (LDA) topic model were generated with pycisTopic v2.0a0 and MALLET v2.0. The SCENIC+ pipeline was run using a precomputed mm10 cisTarget database and motif collection v10nr obtained from the SCENIC+ website, with default parameters except that “mus_musculus” was specified for the species field during motif enrichment. For each eRegulon, we averaged the AUC scores computed from the direct and extended databases.

eRegulons characterizing each cell type and each treatment group within each cluster were identified using the Wilcoxon rank-sum test, as implemented in the Scanpy v1.8.2 sc.tl.rank_genes_groups function, with Bonferroni correction for multiple testing. eRegulon specificity scores (RSS) were calculated from the averaged gene-based AUC scores described above using the regulon_specificity_scores function implemented in SCENIC+ v1.0a1. The top five eRegulons with the highest mean gene-based AUC scores that were detected in more than 10% of cells in each cluster were visualized with dot plots using plotnine v0.12.4. In addition, the expression and RSS values of the top three eRegulons identified as significant cluster-specific markers (adjusted P value < 0.05) were visualized with heat maps using plotnine v0.12.4. eRegulons showing significant differences between stress and control groups were visualized with dot plots using ggplot2 v3.4.4.

*Sox5* and *Sox6* gene expression and eRegulon AUC scores in OPCs from each treatment group were visualized using the VlnPlot function implemented in Seurat v5.0.1. Gene-based eRegulon AUC scores for OPCs were used to construct a nearest-neighbor graph with Scanpy v1.8.2, as described above. This graph was then used to generate UMAP feature plots showing Sox5 and Sox6 eRegulon activity with Scanpy v1.8.2. Sox5 and Sox6 network data, including edges connecting TFs to regions and regions to genes, were extracted from the SCENIC+ object using the create_nx_tables function implemented in SCENIC+ v1.0a1 and plotted with NetworkX v3.2.1. OPC cluster-specific markers and genes differentially expressed between stress and control groups, used for node labeling, were identified with Scanpy v1.8.2 as follows: gene expression counts were normalized with the sc.pp.normalize_total and sc.pp.log1p functions, and differential expression was performed with the sc.tl.rank_genes_groups function, with P values adjusted for multiple comparisons using the Benjamini-Hochberg method. Genes with adjusted P values < 0.05 were selected for labeling in the network plot.

### Isolation of PDGFRα+ OPCs using magnetic-activated cell sorting

OPCs were isolated using magnetic-activated cell sorting (MACS) and anti-PDGFRα MicroBeads (Miltenyi Biotec). In brief, mouse cerebral cortices were dissected, incubated in an enzyme mix, and processed in the gentleMACS™ Octo Dissociator with Heaters for 30 minutes. The resulting suspension was filtered through a 70-μm MACS Smart Strainer. To remove myelin and other unwanted cell types, the cell suspension was centrifuged at 300 × g for 10 minutes at 4°C, and the supernatant was aspirated entirely. The pellet was resuspended in cold D-PBS, mixed with Debris Removal Solution, and overlaid with cold D-PBS. The top two phases were removed after centrifugation at 3,000 × g for 10 minutes at 4°C. The pellet was washed in cold D-PBS and centrifuged at 1,000 × g for 10 minutes at 4°C. Cells were resuspended and incubated with anti-PDGFRα MicroBeads (Miltenyi Biotec) for 15 minutes at 4°C. The labeled cells were then passed through an MS column placed in a magnetic separator. The PDGFRα-positive cells were retained in the column. After several wash steps to remove non-specifically bound cells, the column was removed from the magnetic field, and the OPC-enriched fraction was eluted in MACS buffer. Finally, the positive PDGFRα cells were plated in 24-well plates for subsequent experiments. Tissue dissection, cell sorting, and plating were conducted under sterile conditions.

### Adult OPC culture, differentiation and migration

As described above, the PDGFRα+ cells were isolated from the cortices of 4.5-month-old adult mice using MACS technology. The isolated cells were resuspended in OPC proliferation medium consisting of 50 μL Neurobasal-A Medium (Gibco), 2% MACS NeuroBrew-21, 1% Normocin, 0.5 mM L-glutamine, 10 ng/mL Human PDGF-AA, 10 ng/mL Human FGF-2, and 4 μg/mL forskolin. Cells were then plated at a density of 25,000 cells/cm² on a 24-well plate coated with 0.01% poly-L-lysine and 10 µg/mL laminin. Half of the medium was changed every other day until use. To induce OPC differentiation, the culture medium was replaced on DIV4 with OPC differentiation medium supplemented with 40 ng/mL T3 and 10 ng/mL CNTF. Every 2-3 days, half of the medium was replaced with fresh differentiation medium until analysis.

To assess OPC migration, OPCs maintained in proliferation medium were analyzed on DIV6. Time-lapse images were acquired over 24 hours using the ImageXpress Micro system (Molecular Devices) and MetaXpress software. Cell migration was analyzed in ImageJ, with single-cell trajectories tracked manually using the Manual Tracking plugin or automatically using TrackMate. Migration parameters, including speed, total distance, velocity, and displacement, were quantified and analyzed using GraphPad Prism.

### Primary OPC culture

OPCs were isolated from the cerebral cortices of postnatal (P0-P1) mice using MACS (Miltenyi Biotec) according to the manufacturer’s instructions. The purified OPCs were then resuspended in an OPC culture medium, which consisted of neurobasal medium supplemented with 2% MACS NeuroBrew-21, 1% penicillin-streptomycin, 0.5 mM L-glutamine, 10 ng/mL human PDGF-AA, and 10 ng/mL human FGF-2. Cells were plated onto poly-D-lysine-coated culture plates at a density of 10,000 cells/cm² and incubated at 37°C in 5% CO₂. The culture medium was refreshed every 2-3 days, and cell viability and proliferation were regularly monitored.

### Immunocytochemistry

Cells were fixed with 4% paraformaldehyde for 10 minutes and then washed three times with PBS for 5 minutes at room temperature (RT). Permeabilization was done using 0.1% Triton X-100 in PBS (PBST) for 10 minutes, followed by blocking non-specific binding with 5% goat serum in PBST for 1 hour. Primary antibodies were applied overnight at 4 °C, including Olig2 (Synaptic Systems, #292015) (1:100), PDGFRα (Abcam, #ab203491) (1:400), and MBP (Abcam, #ab218011; 1:400). The next day, coverslips were washed three times with PBS for 5 minutes each and incubated with secondary antibodies for 1 hour at RT. After two additional 5-minute washes with PBS, DAPI Fluoromount-G (Southern Biotech) was applied for nuclear staining and mounting. Images were acquired using Echo microscopy at 20× magnification, with a scale bar of 90 μm. All 2D image analyses were performed using Fiji (version 1.53c).

### Immunofluorescence staining of brain sections

Mice were perfused with cold PBS, followed by 4% paraformaldehyde in phosphate buffer, and processed for OCT embedding. Tissue sections (15 μm) were washed, blocked with 5% serum in TBST for 1 hour, and incubated overnight at 4°C with the appropriate primary antibody (Supp. Table 22). Subsets of sections were stained with antibodies targeting synaptic proteins and glutamate receptors. After washing, sections were incubated with Alexa-conjugated secondary antibodies (1:500) for 1 hour at room temperature, followed by mounting in ProLong™ Gold with DAPI. Images were captured at 20x magnification using a Nikon N-Storm microscope, with additional images obtained at 40x or 100x using a Nikon STORM microscope with an oil lens.

### Image analysis

ImageJ was used to quantify synaptic counts and cell intensity. Images were converted to grayscale, and a threshold was applied to distinguish synaptic puncta and cells from the background. The ‘Analyze Particles’ function was used to count puncta or cells, applying size and circularity filters to exclude noise. Puncta density (puncta/µm²) was calculated for markers such as Bassoon and Homer. In some cases, GluD1 and GluK2 intensities were measured in PDGFRα-positive cells by applying a threshold to select these cells and using the ‘Analyzè > ‘Measurè function. Intensity values were normalized to control levels, and fold change in intensity was calculated by comparing experimental groups to controls.

For fractal analysis, fractal dimension (DB) was quantified using the FracLac plugin for ImageJ, which employs a box-counting method to determine the level of detail in cell contours at increasing scales. The fractal dimension was calculated using the formula DB = Log(N)/Log(r), where N is the number of pixels at a given scale factor (r). Before analysis, the images were converted to grayscale. Subsequently, a threshold was applied to transform the grayscale image into a binary format, and a region of interest was selected to improve measurement accuracy. Statistical analysis was conducted on three different biological replicates of three independent cultures, with data from 40 cells per condition. The statistical significance between the stress and control groups was determined using a *t*-test with a significance level set at *p* < 0.05.

### Lentivirus production and transduction

Lentiviral vectors were produced as previously described ^67^. Briefly, 18 μg of the lentiviral transfer vector was co-transfected with 18 μg of psPAX2 (Addgene #12260) and 6 μg of pMD2.G (Addgene #12259) into 293FT cells (Clontech) grown in a 150 mm dish using polyethylenimine (PEI, Polysciences) as the transfection reagent. Eight hours post-transfection, the medium was replaced with fresh culture medium. Forty-eight hours later, viral supernatants were harvested, cleared by centrifugation (2000 rpm, 10 min), and filtered through a 0.45 μm PES filter. Viral particles were then pelleted by ultracentrifugation at 20,000 rpm for 2 hours using an Optima XE100 ultracentrifuge (Beckman Coulter) and resuspended in 50–100 μL of PBS. Packaged lentiviruses were diluted in OPC culture medium and applied to OPCs in the presence of 4 μg/mL polybrene. Twenty-four hours later, the medium was replaced with fresh culture medium. Ultrapure lentiviral products for in vivo injection were packaged by the NeuroTools Core facility in University of North Carolina at Chapel Hill.

### Stereotaxic lentivirus injection

Mice were stereotaxically injected bilaterally with lentiviral vectors expressing either GFP or Sox6-T2A-GFP into the dorsal hippocampus. Identical viral doses were used for both vectors, with 3×10^7^ IU delivered in 0.5 µL per hemisphere. Injections were performed at coordinates corresponding to bregma -2.1 to -2.3 mm. Viral infusions were delivered using a microinjection pump (Dual Syringe, Model 11, Harvard Apparatus, MA-70-2209) at a rate of 0.05 µL/min to a final volume of 0.5 µL per hemisphere. Infusions were administered over 10 min, followed by an additional 5 min to allow diffusion before needle withdrawal and minimize backflow. The total injection time per site was approximately 15 min. Mice were allowed to recover for three weeks before being subjected to the CUMS or control procedure.

### Quantitative reverse transcription-polymerase chain reaction (qRT-PCR)

Brain RNA was extracted using a Direct-zol RNA Miniprep kit in the presence of RNase-Free DNase (Zymo Research, TX, USA), following the manufacturer’s instructions. RNA integrity was assessed and quantified using NanoDrop. cDNA was generated using a High-Capacity cDNA Reverse Transcription Kit (Applied Biosystems). Real-time polymerase chain reaction was performed using TaqMan Fast Advanced Master Mix (Applied Biosystems, #4444557), along with the appropriate TaqMan probes: Cnp (Mm01306641_m1), Mbp (Mm01266402_m1), and Plp1 (Mm01297210_m1). Mouse glyceraldehyde 3-phosphate dehydrogenase (GAPDH) was used as the internal control to standardize the samples. PCR was performed under the following conditions: 50 °C for 2 minutes, followed by 95 °C for 10 minutes, and then 40 cycles of 95 °C for 15 seconds and 60 °C for 1 minute. Each sample, derived from three biological replicates (n = 3-4 animals per condition) and technical replicates (triplicates per sample), was analyzed using a CFX384 Real-Time PCR system (Bio-Rad). Negative controls, including no-template controls, were included to assess contamination or non-specific amplification. Data analysis was performed using the comparative CT method (2−ΔΔCT). Statistical significance was determined using GraphPad Prism, applying either the t-test or ANOVA, with a significance threshold set at *p* < 0.05.

### ChIP-seq, ChIP-qPCR, and Chromatin accessibility tests

ChIP was performed using the SimpleChIP Plus Enzymatic Chromatin IP Kit (cat. no. 9003, Cell Signaling Technology) according to the manufacturer’s instructions. Two biological replicates were included. For each replicate, 4×10⁶ OPCs were isolated from P2-P4 mice and fixed with 1% formaldehyde to crosslink histone and non-histone proteins to DNA. Chromatin was digested with micrococcal nuclease into 150–900 bp DNA-protein fragments, corresponding to approximately one to five nucleosomes. Two percent of the chromatin sample was reserved as input control, and the remaining chromatin was subjected to ChIP using an anti-SOX6 antibody (Abcam, ab30455, 2 μg/sample). DNA library preparation and sequencing were performed by the Vanderbilt Technologies for Advanced Genomics core facility. Paired-end sequencing was performed on a NovaSeq XP platform with 150-bp reads, targeting an average of 40 million reads per sample.

ChIP-seq reads were processed with the nf-core/chipseq pipeline v2.1.0 using default settings for read preprocessing and alignment to the mouse mm10 reference genome ^68^. The mm10 blacklist was used to exclude problematic genomic regions, and aligned BAM files were generated for downstream peak calling ^69^. ChIP-seq signal was summarized around transcription start sites (TSS; ±20 kb) using deepTools computeMatrix, and visualized as a heatmap with deepTools plotHeatmap ^70^. Aligned reads from ChIP and matched input replicates were combined into HOMER tag directories using makeTagDirectory ^53^. Peaks were then called with HOMER findPeaks in TF mode and the resulting peak list was filtered to retain only UCSC-style chromosomes prior to downstream analyses ^53^. Peak coordinates were annotated to genomic features and nearest genes using HOMER annotatePeaks.pl against the mm10 reference genome. Enrichment of JASPAR2024 Mus musculus CORE TF motifs was evaluated with monaLisa v1.16.0 calcBinnedMotifEnrR function using the mm10 genome as background and an oversampling parameter of 4 ^71, 72^. Top 10 most significant motifs were visualized with monaLisa v1.16.0 plotMotifHeatmap. De novo motif discovery was performed with HOMER findMotifsGenome.pl with the mm10 genome reference using default parameters and masking repetitive sequences ^53^.

For ChIP-qPCR analysis, ChIP DNA was prepared using the Low Cell ChIP Kit (cat. no. 53086, Active Motif). ChIP was performed using six biological replicates. For each replicate, 30,000 OPCs isolated from neonatal mice were fixed as described above and sonicated using a Bioruptor 300 sonicator. The fragmented chromatin was divided equally for ChIP using preimmune IgG (Cell Signaling Technology, #2729), anti-SOX6 (Abcam, ab30455), or anti-H3K27ac (Abcam, ab4729) antibodies. Purified ChIP DNA was then analyzed by qPCR using the following primers: *Pik3r1* peak, F: 5′-GTTTGAAGAAGGCCTCTCCTTTG-3′, R: 5′-GTCTCTTTTGTACGGAGCACA-3′; *Ntn1* peak, F: 5′-GGCGGCGATTTATTCTGGGA-3′, R: 5′-ACCCCGGGCTAATCTGAACA-3′; *Nckap5* peak, F: 5′-TCTCTGGTTGGAGGGTAGCG-3′, R: 5′-ATCCCGATCACGAGGAGAAC-3′; *Rgcc* peak 1, F: 5′-GTGGCCCAGTGGTTCTTTCTC-3′, R: 5′-GACCTTCTGCGGTTTCGGG-3′; and *Rgcc* peak 2, F: 5′-TTCTATGTGGAGACACGTAGTGG-3′, R: 5′-ACAAAAGAAGGCTACCCCCTC-3′.

Chromatin accessibility was assessed using the EpiQuik Chromatin Accessibility Assay Kit (cat. no. P-1047, EpigenTek) according to the manufacturer’s protocol. For each reaction, 0.2×10⁶ OPCs isolated from control or stressed mice were used, with six biological replicates per group. Briefly, chromatin was isolated from each sample and divided into two equal fractions. One fraction was treated with the nuclease mix, whereas the other was retained as an undigested control. DNA was then purified and analyzed by qPCR to assess chromatin accessibility at selected genomic regions using primers described above.

### Language editing

During manuscript preparation, the authors used ChatGPT-4o for language editing, specifically to correct grammatical errors and improve clarity. The use of ChatGPT was strictly limited to language editing. After using this tool, the authors reviewed and revised the manuscript as needed and take full responsibility for the content of the publication.

## Acknowledgements

The authors thank Destany Ware for technical support. We also thank members of the Jiao and Wang labs for their input.

## Author contributions

Conceptualization: Q.W., and K.J.; Methodology: A.B., C.M.J., Y.S., S.S., H.W., W.Z., H.S., Q.W. and K.J.; Data analysis: A.B., K.J., C.M.J., Y.S., S.S., Y.G.G., T.J.L. and Q.W.; Investigation: A.B., C.M.J., Y.S., S.S., W.L., Y.G.G., E.D., Q.W. and K.J.; Resources: Q.D., A.S., H.S., W.Z., Q.W. and K.J.; Manuscript preparation: A.B., C.M.J., Y.S., S.S., K.J. and Q.W.; Supervision: K.J. and Q.W.; Funding acquisition: K.J. and Q.W.

## Data and material availability

All data needed to evaluate the conclusions in the paper are present in the paper and/or the Supplementary Materials. Original sequencing reads have been deposited in the Gene Expression Omnibus (GEO) under accession numbers GSE330734 and GSE330735.

