## Supplementary material for "Chronic Mild Stress Impairs Hippocampal Myelination through SOX6-Dependent Dysfunction of Oligodendrocyte Lineage Cells": Supp Figures

### Supplementary Information

#### Supplementary Figures

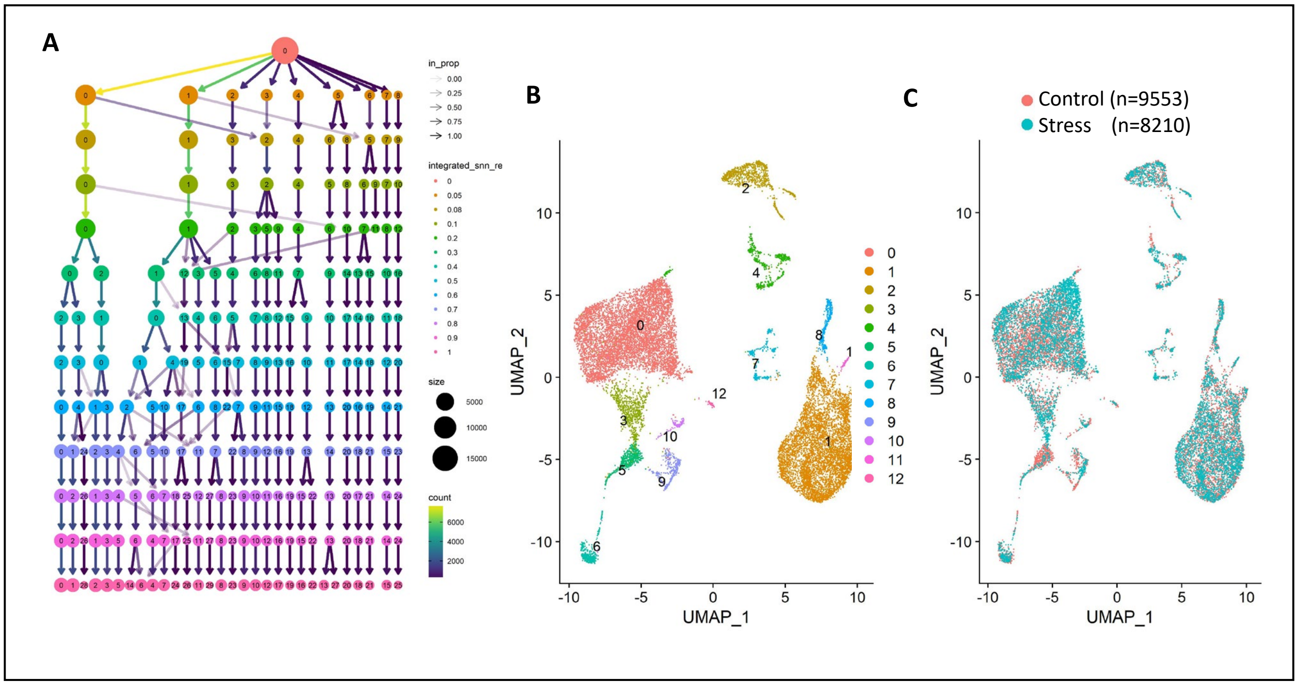


**Sup. Fig. S1. (A)** The Clustree plot shows how clusters change across different resolutions. **(B)** The UMAP chart shows that nuclei isolated from the hippocampus can be grouped into 13 clusters. **(C)** This chart displays the distribution of nuclei from control (red) and stress (blue) groups across all clusters.

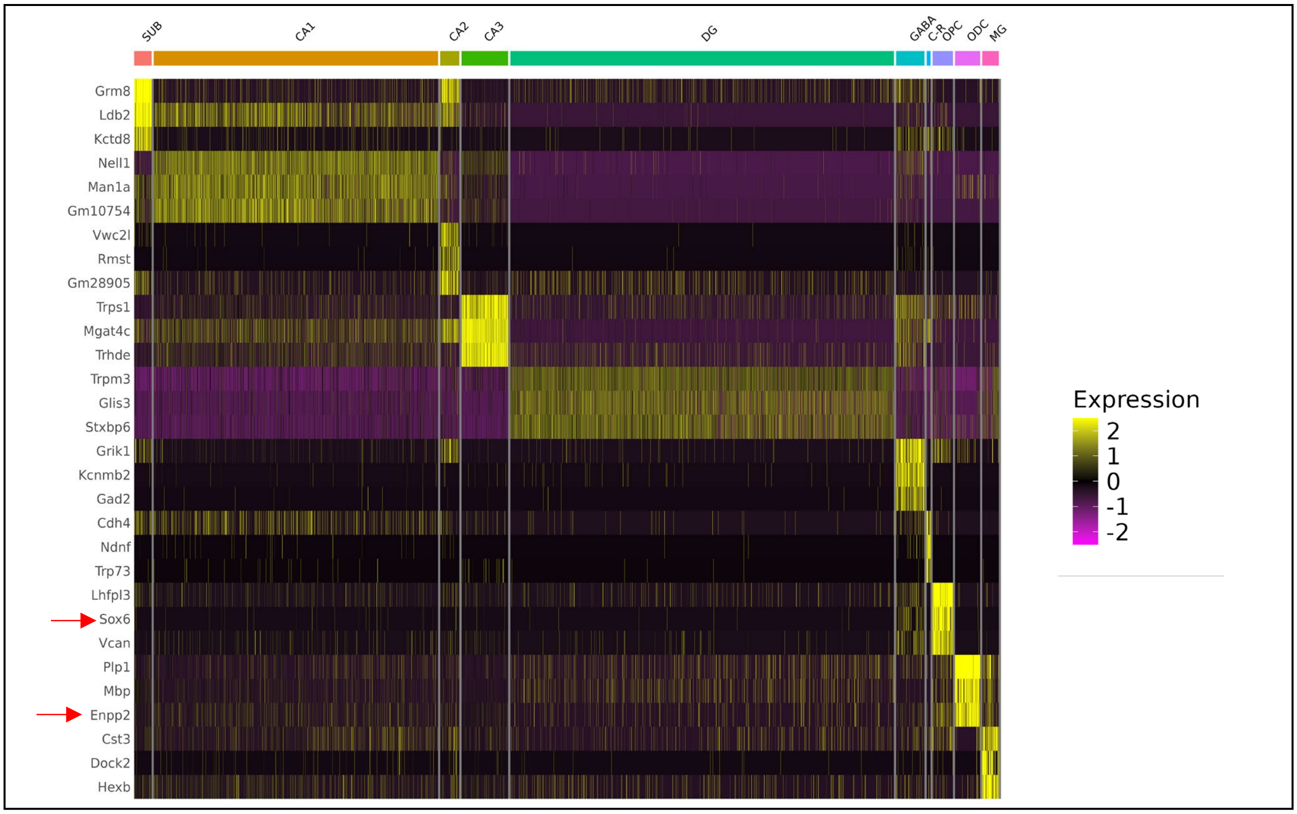


**Sup. Fig. S2.** The heatmap displays the top three positive marker genes for each cluster. SUB: subiculum neurons; CA1-3: neurons in CA1-3 regions; DG: dentate gyrus neurons; GABA: GABAergic neurons; C-R: Cajal-Retzius neurons; OPC: oligodendrocyte progenitor cells; ODC: oligodendrocyte cells; MG: microglia.

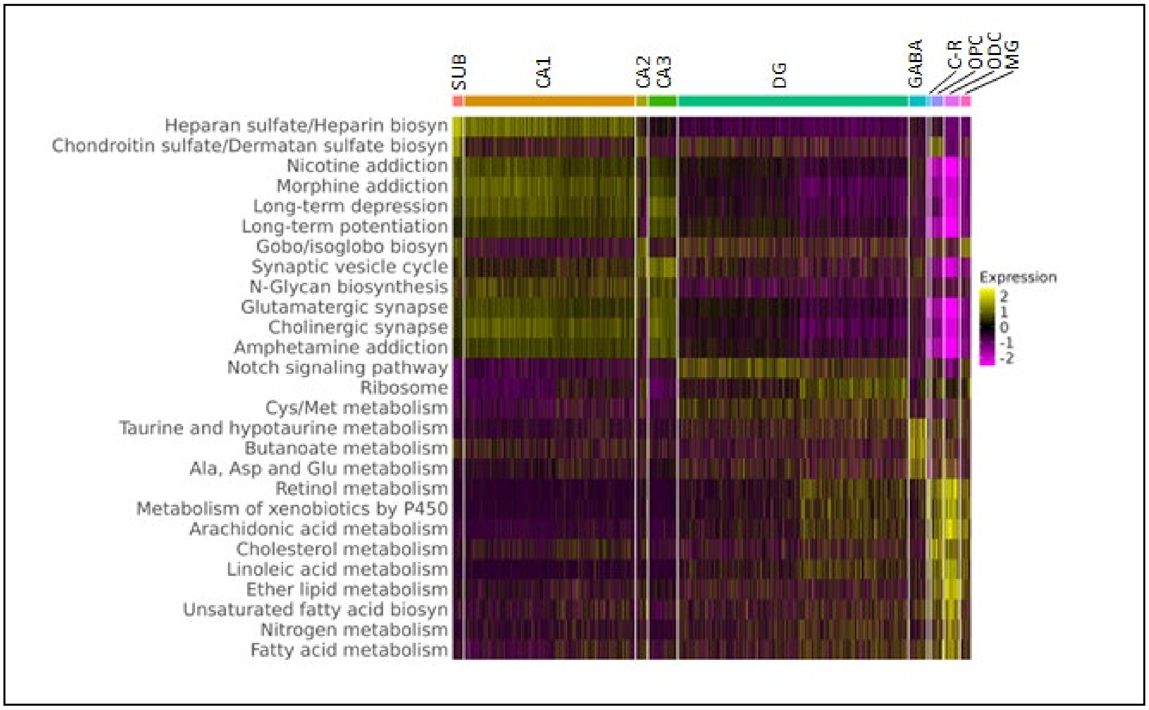


**Sup. Fig. S3.** The gene expression profiles of each cluster were analyzed using AUCell with KEGG gene sets and terms. Three examples of the top enriched KEGG terms associated with each cluster are shown in the heatmap. SUB: subiculum neurons; CA1-3: neurons in CA1-3 regions; DG: dentate gyrus neurons; GABA: GABAergic neurons; C-R: Cajal-Retzius neurons; OPC: oligodendrocyte progenitor cells; ODC: oligodendrocyte cells; MG: microglia.

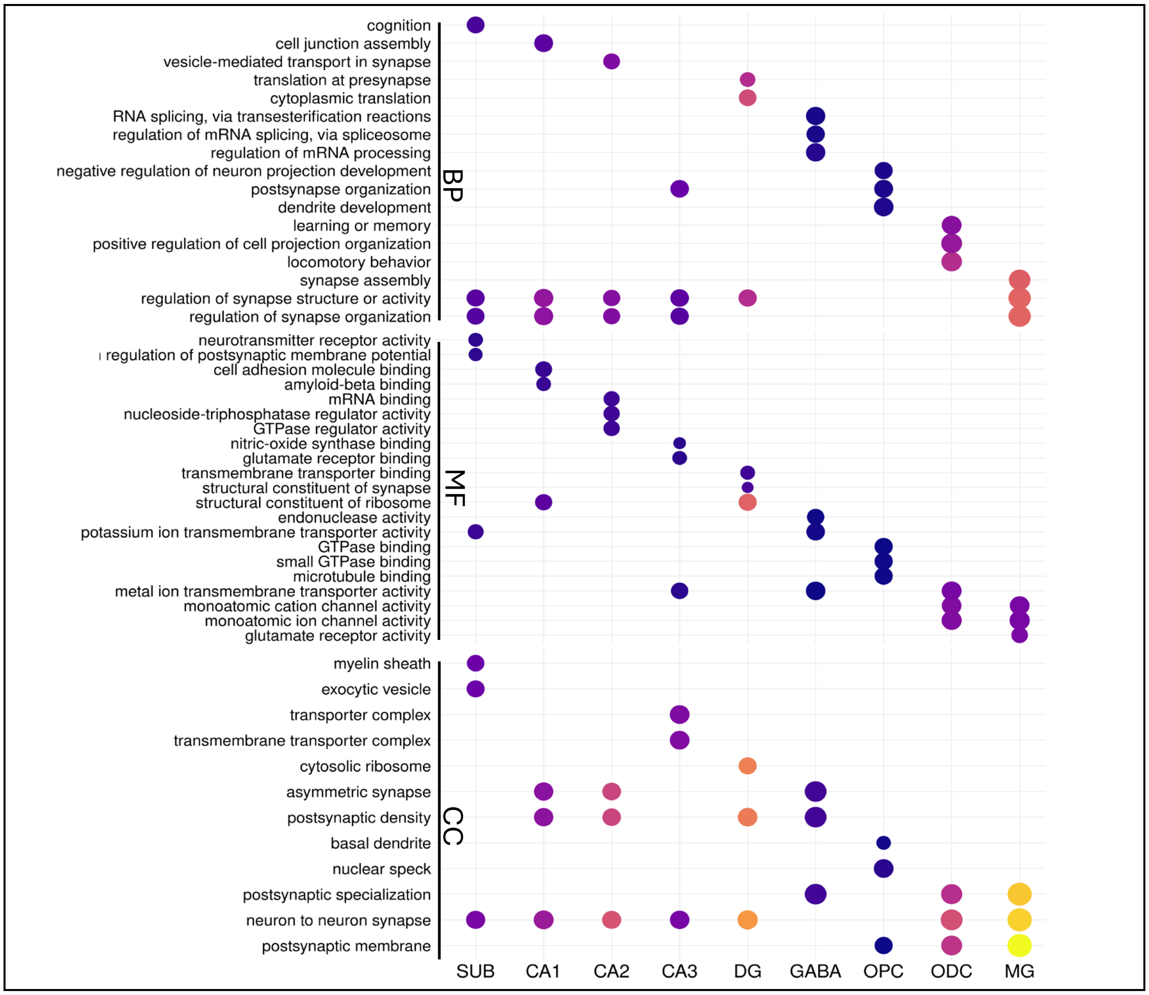


**Sup. Fig. S4.** The dot plot shows the top three significantly enriched GO terms among DEGs for each cell population. BP: Biological Process; MF: Molecular Function; CC: Cellular Component.

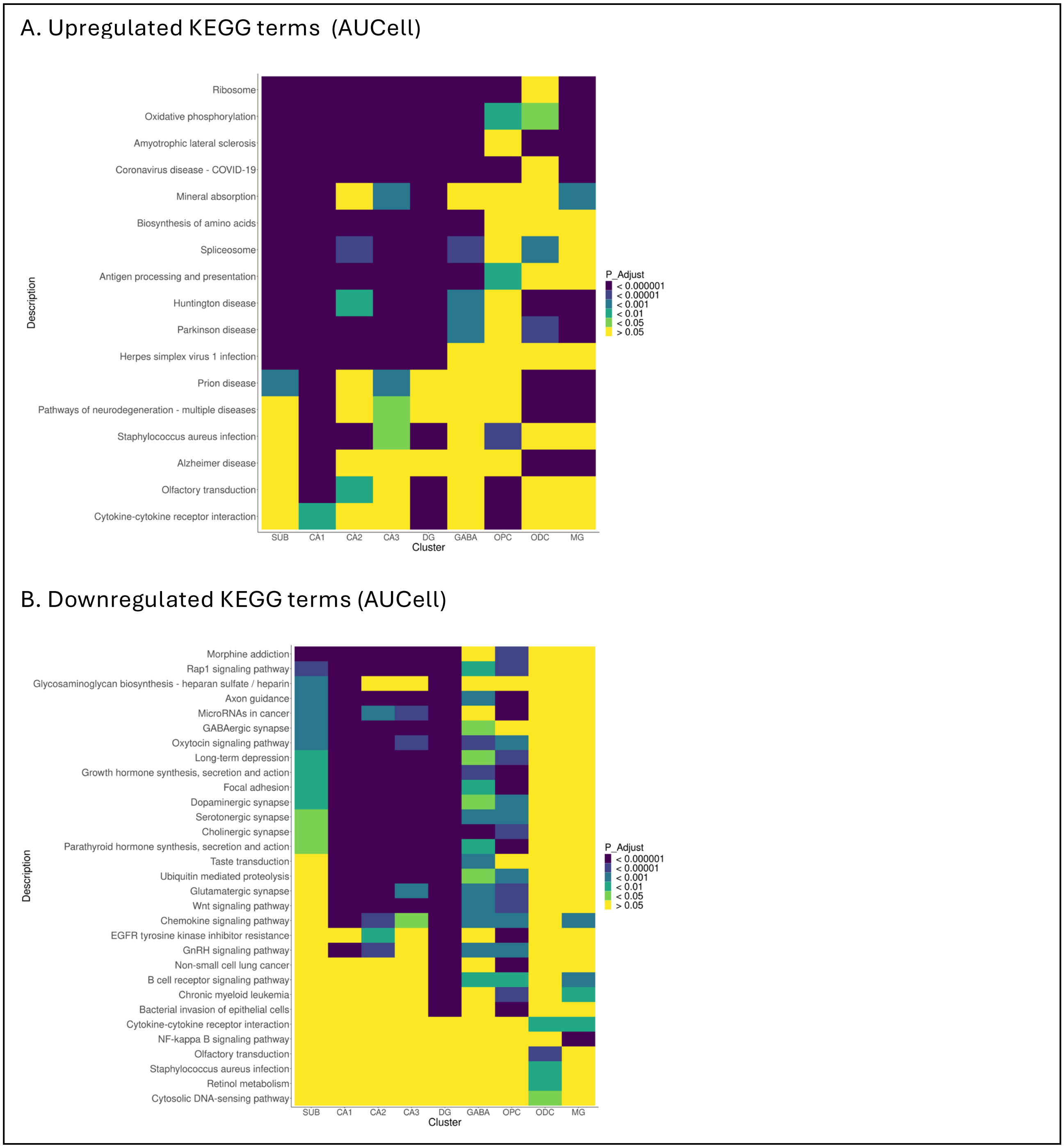


**Sup. Fig. S5.** The top three upregulated **(A)** and downregulated **(B)** terms for each cluster from AUCell analysis using KEGG gene sets and terms.

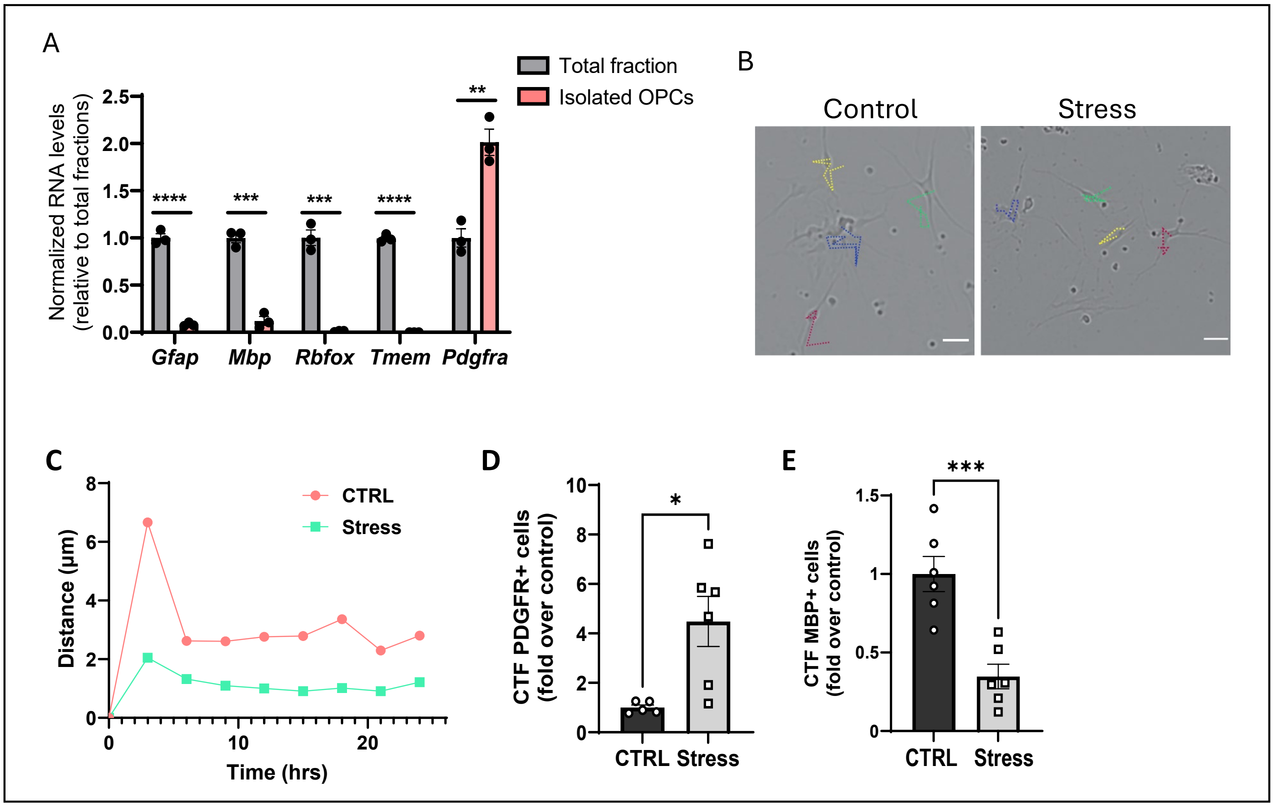


**Sup. Fig. S6.** Adult OPC culture, migration, and differentiation. **(A)** Adult OPCs were isolated from control or stressed mice following procedures described in Material and Method. Relative expressions of maker genes for different cell types in isolated OPCs were examined by qRT-PCR and presented as fold change versus the level in the total fraction. **, $p$<0.01; ***, $p$<0.001; ****, $p$<0.0001 by unpaired t-test. Data are mean ± SEM. **(B)** Spontaneous migration was monitored on DIV5 for 24 hrs. Representative images of OPC culture on DIV6. The dotted line represents the migration trajectory in 24 hrs. **(C)** The quantitation of the average distance migrated over a 24-hour period. N = 35–40 cells from three independent cultures per group. **(D,E)** In a separate set of cultures, OPC differentiation was induced on DIV 4, and the cells were subjected to immunofluorescence staining on DIV 7. Quantifications of corrected total fluorescence (CTF) in PDGFα$^{+}$ cells (D) and MBP+ cells (E) are shown.

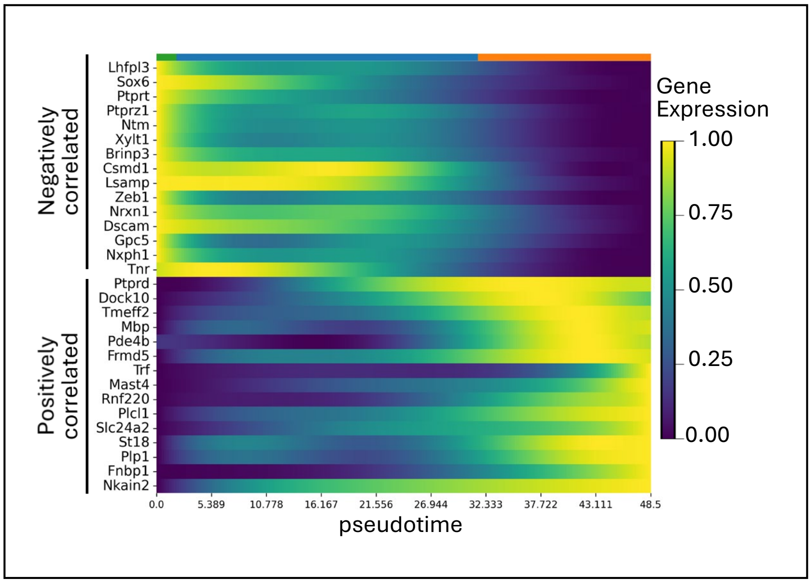


**Sup. Fig. S7.** Top 15 genes whose expression is positively and negatively correlated with pseudotime**.**

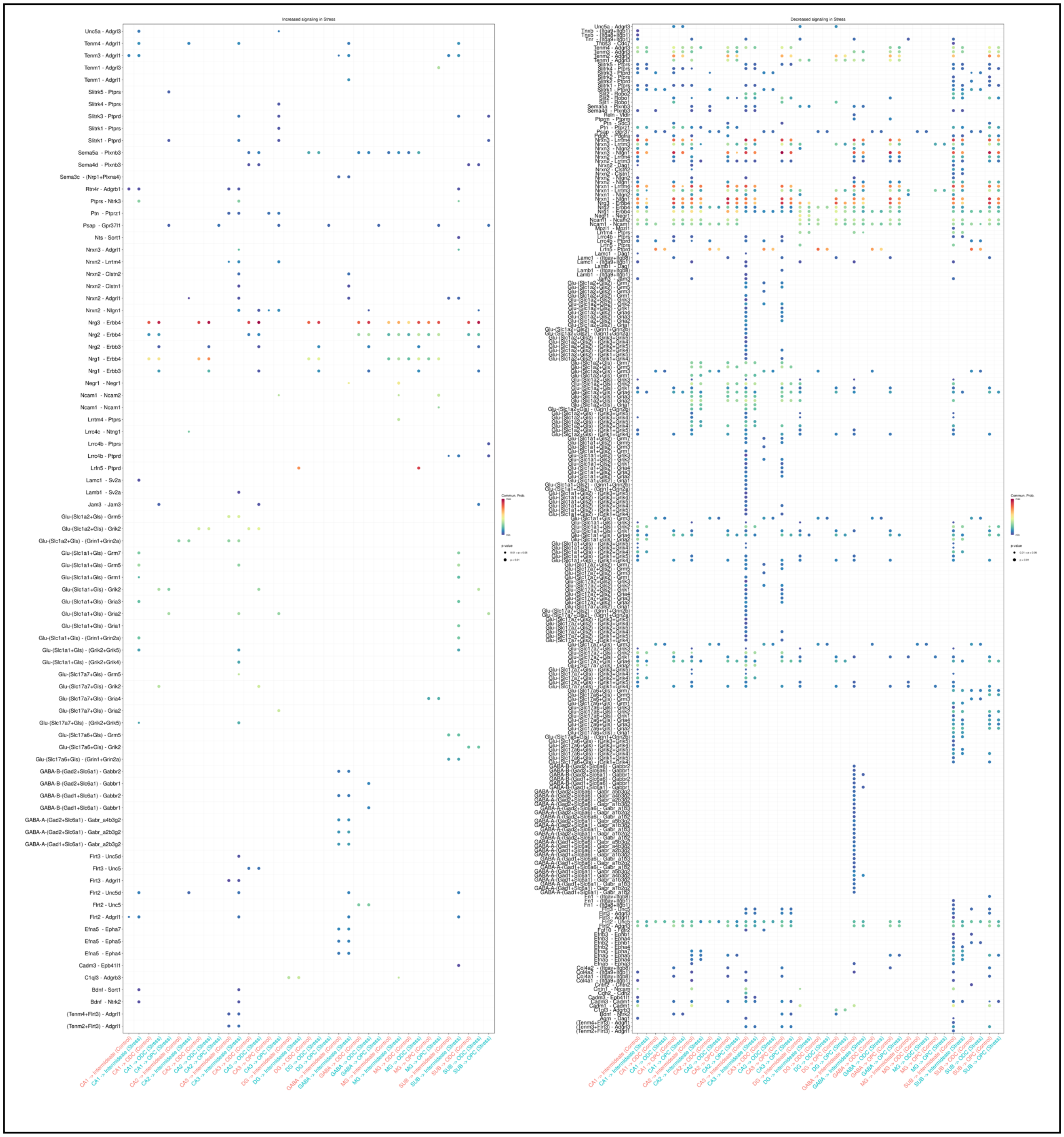


**Sup. Fig. S8.** This dot plot summarizes all significantly altered cell-cell communication pathways incoming to OPC-m3, Intermediate, and ODC-m3 under stress.

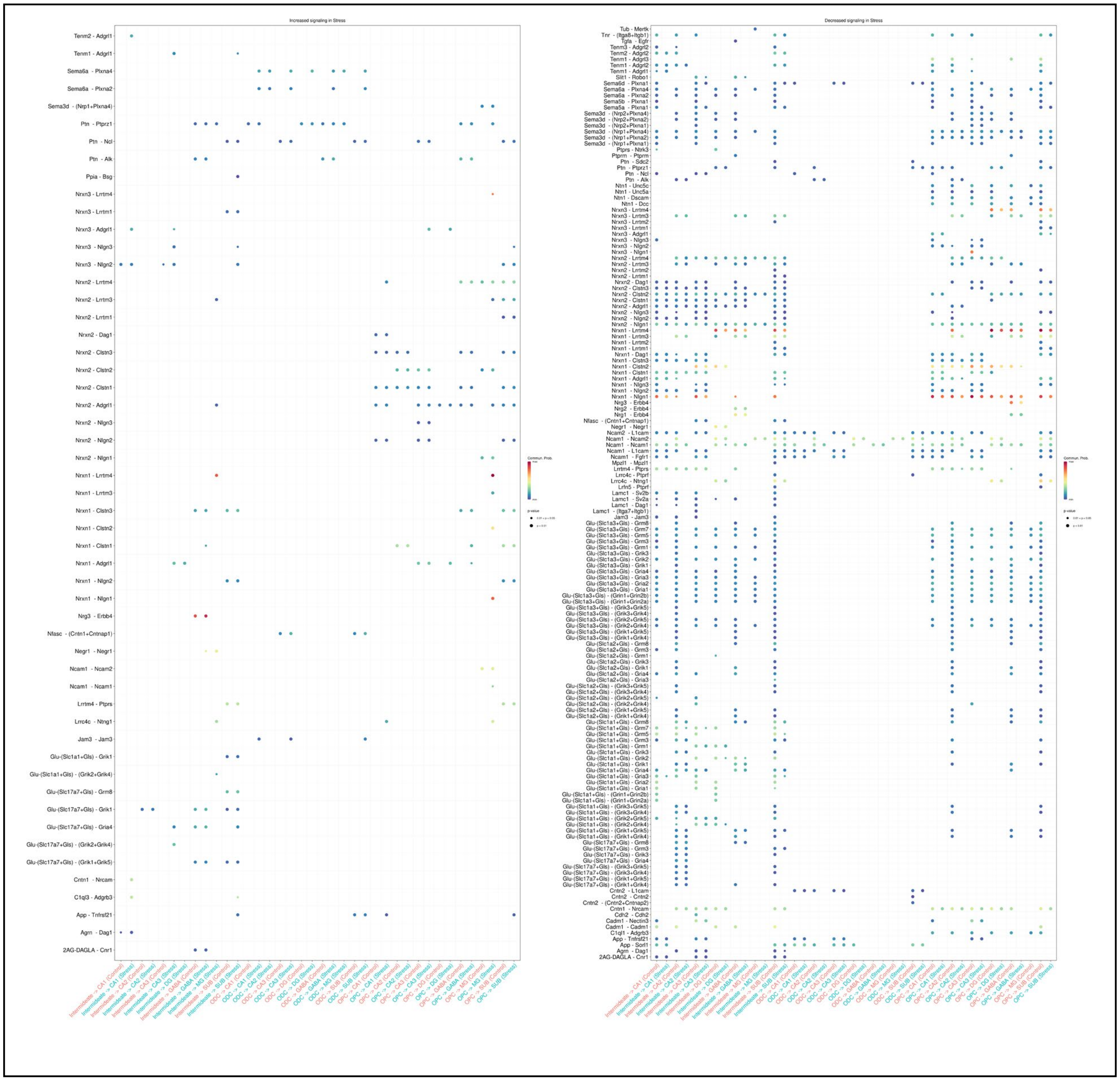


**Sup. Fig. S9.** This dot blot summarizes all significantly altered cell-cell communication pathways outgoing from OPC-m3, Intermediate, and ODC-m3 under stress.

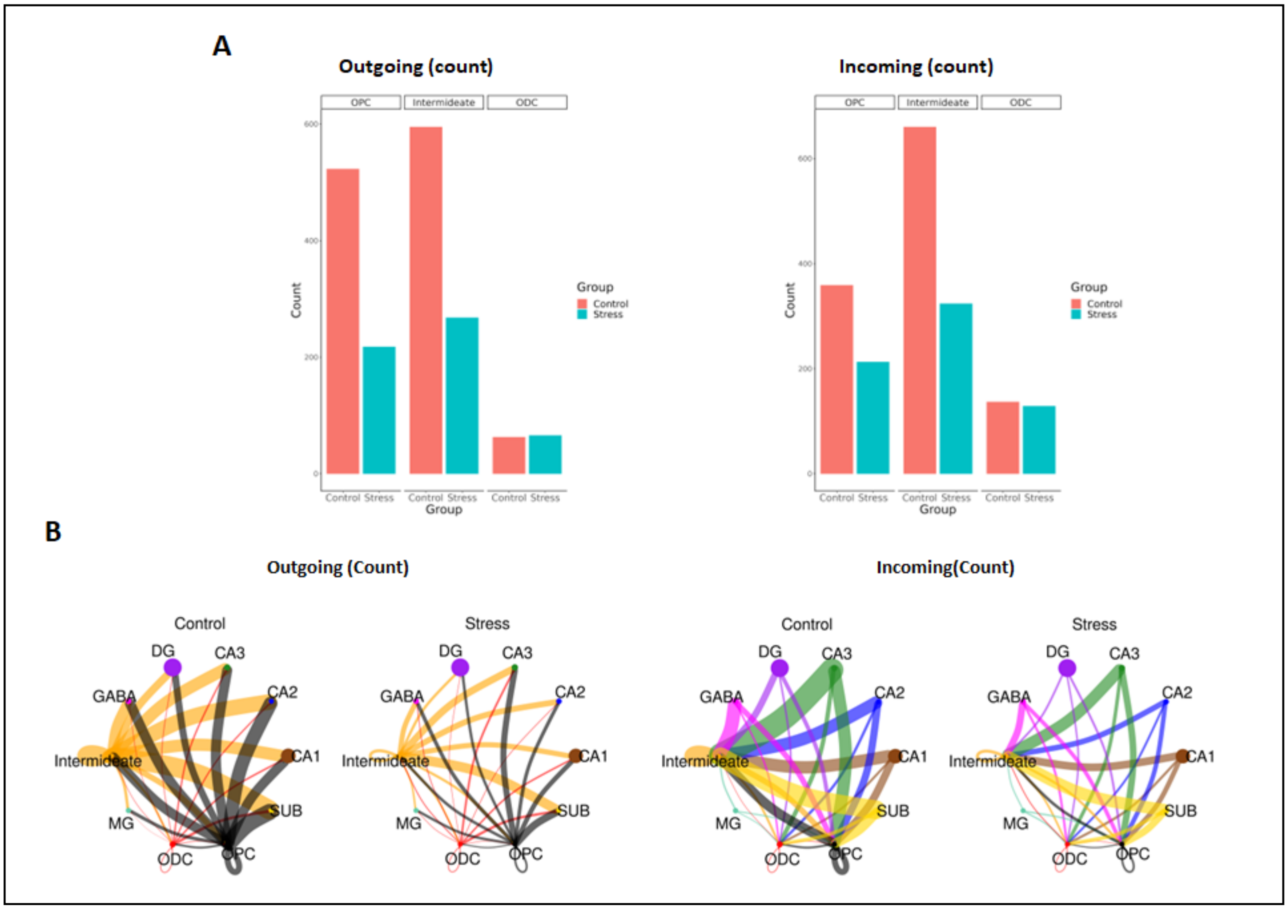


**Sup. Fig. S10. (A)** The bar charts summarize the counts of outgoing (left panel) and incoming (right panel) intercellular communication pathways associated with OPC-m3, Intermediate, and ODC-m3 using CellChat analysis. **(B)** The circle plots display the counts of outgoing (left panels) and incoming (right panels) communication networks of OPC-m3, Intermediate, and ODC-m3 with other clusters under both control and stress conditions.

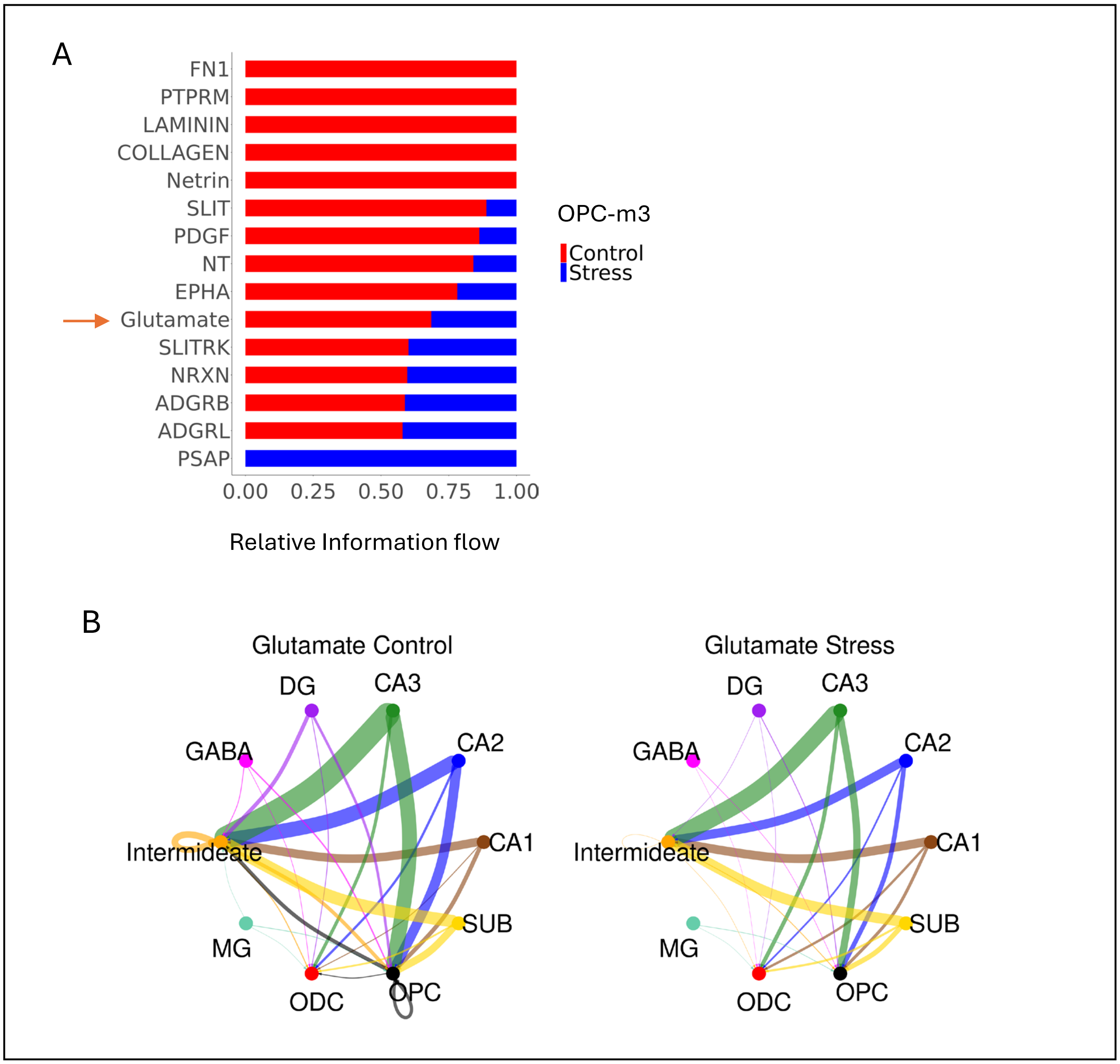


**Sup. Fig. S11.** **(A)** The bar chart displays the incoming signaling pathways to OPC-m3 with significantly altered relative information flow under stress, as determined by CellChat. Glutamate signaling is indicated with a red arrow. **(B)** The circle plots illustrate the incoming glutamate signaling probability in OPC-m3, Intermediate, and ODC-m3 cells from all clusters.

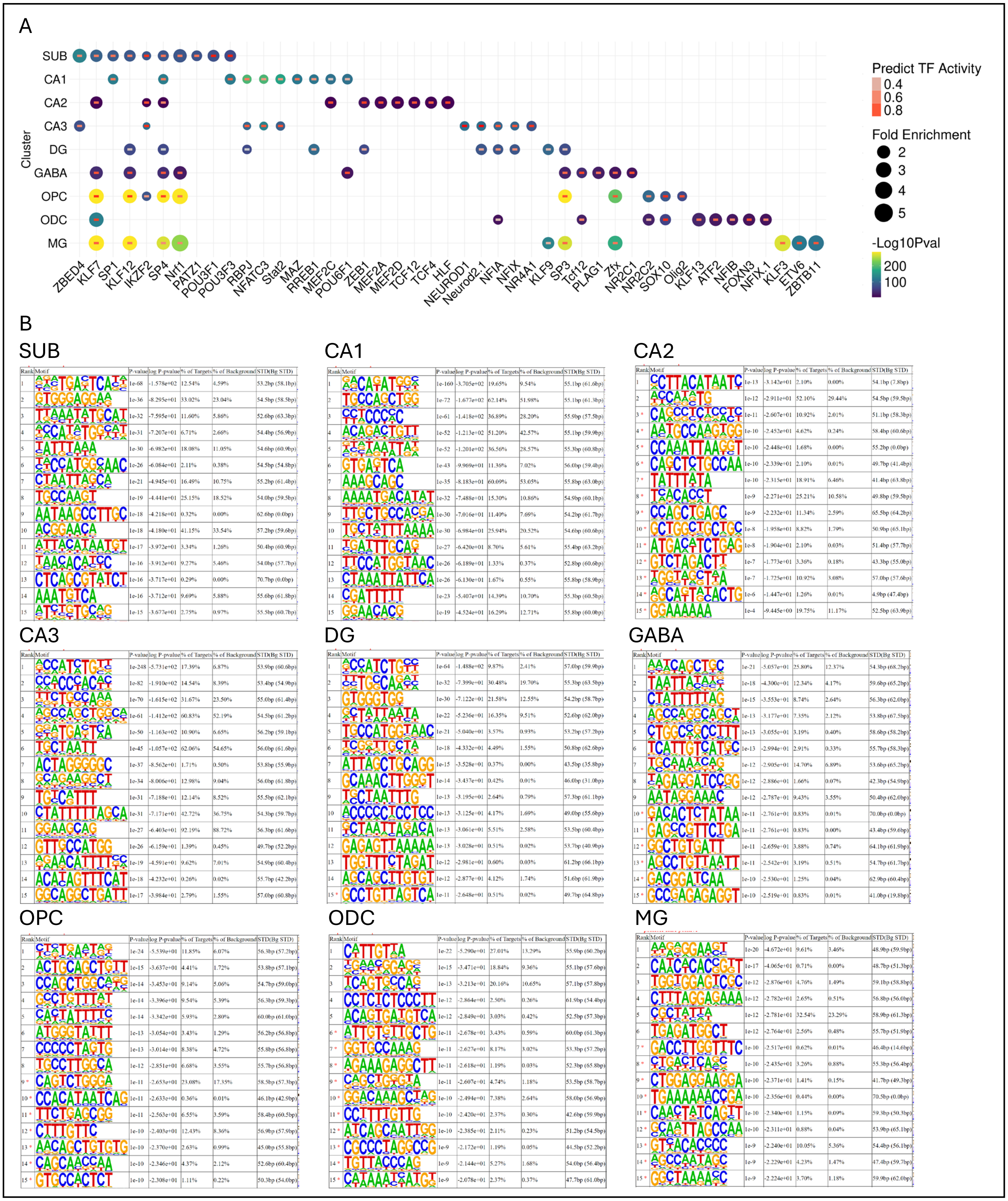


**Sup. Fig. S12. (A)** The DA peaks of each cluster, located within 100 kb of a cluster marker gene and significantly correlated with marker gene expression, were analyzed using JASPAR. The top

10 significantly enriched TF binding motifs in each cluster are shown in the dot plot. **(B)** The top 15 *de novo* motifs within the DA peaks for each cluster were identified using HOMER.

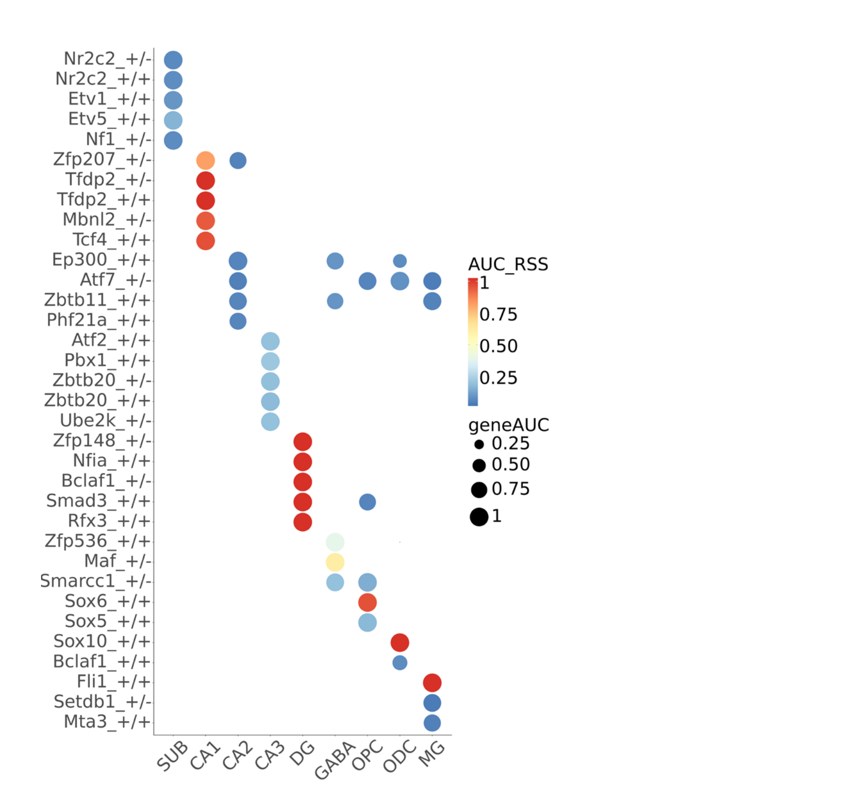


**Sup. Fig. S13.** Key transcriptional regulatory network units identified through SCENIC+ analysis. The dot plot displays the top five eRegulons with the highest mean AUC values in each cluster. AUC_RSS refers to the eRegulon Specificity Score, calculated using AUC values from transcriptome expression profiles (geneAUC), which measures the specificity of an eRegulon within a given cluster.

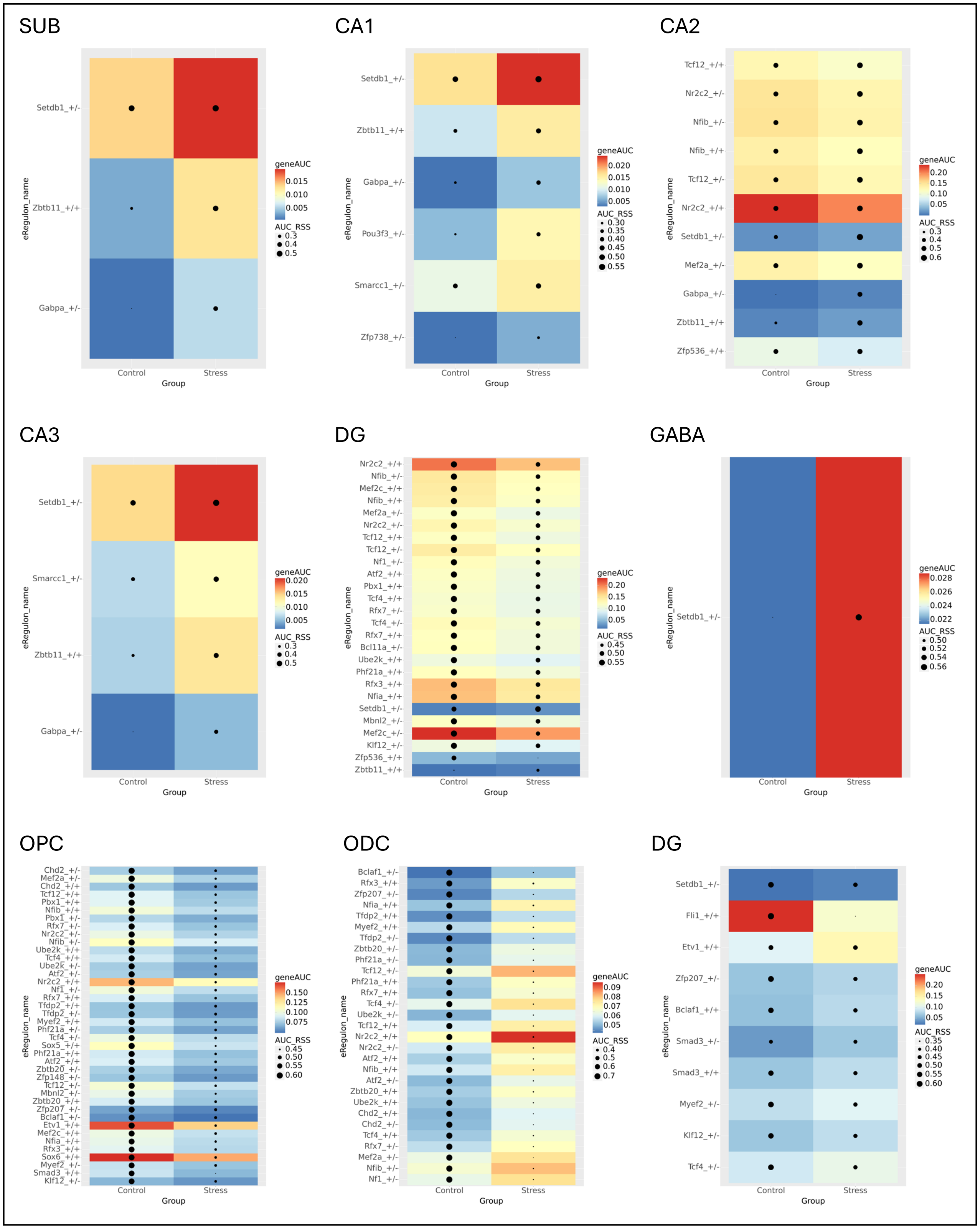


**Sup. Fig. S14.** The heatmaps display eRegulons with significantly altered activity under stress in each cluster.

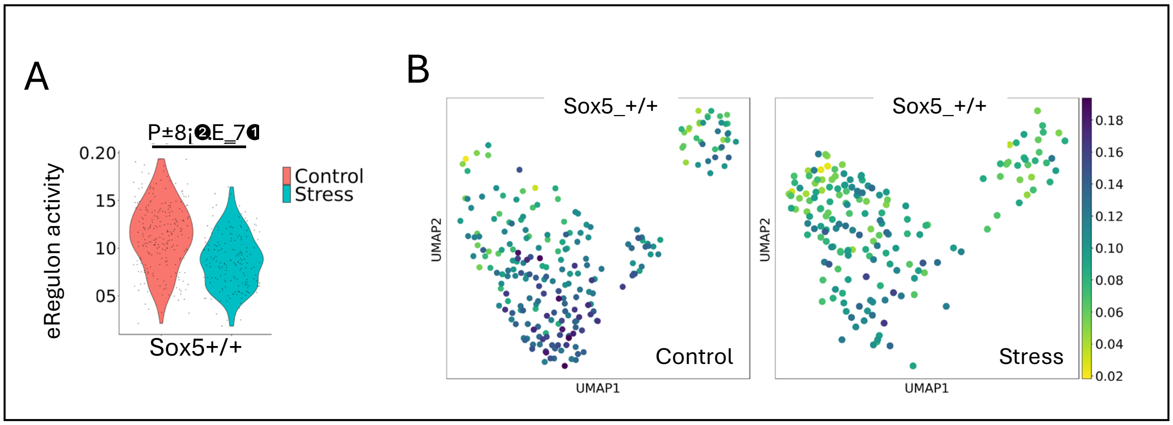


**Sup. Fig. S15.** SOX5 eRegulon activity is reduced by CUMS. **(A)** SOX5 eRegulon activity in control versus stress groups. **(B)** OPCs were re-clustered based on eRegulon activity. Feature maps illustrate the decreased activity of *Sox5*-mediated eRegulons under stress at the single-nucleus level.

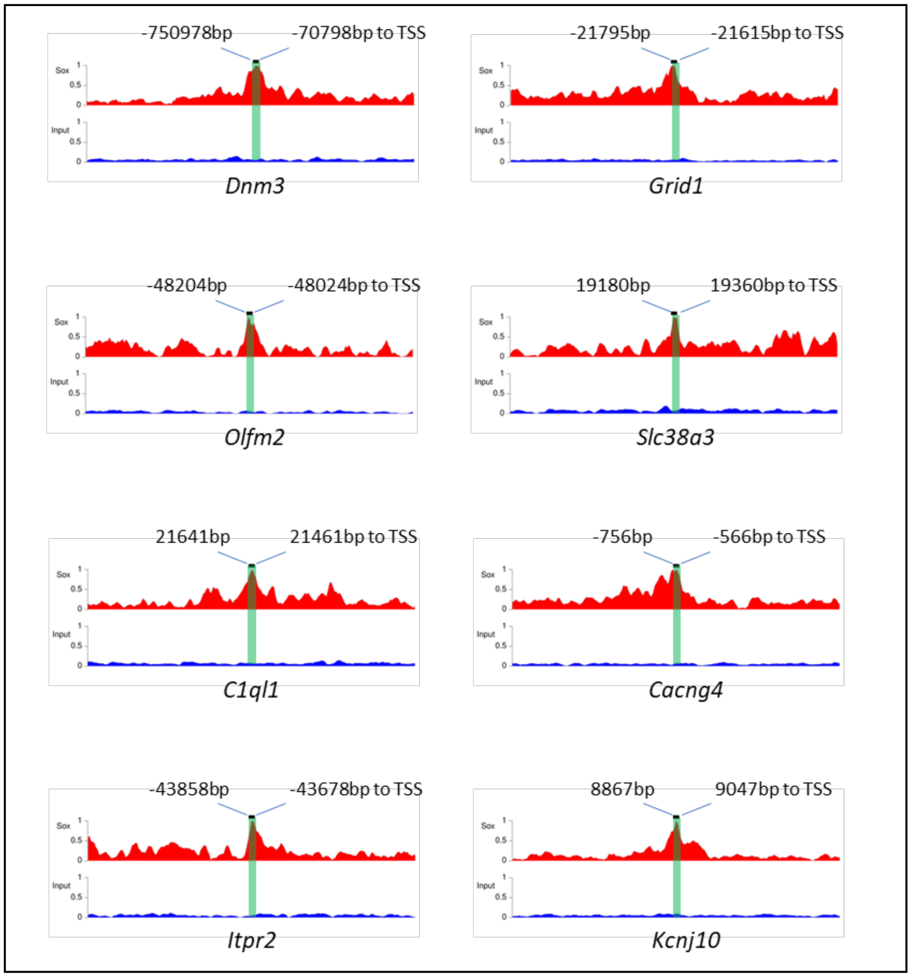


**Sup. Fig. S16.** Coverage plots showing the peaks associated with the eight genes that are known to regulate the glutamatergic signaling. These peaks are with 100kb upstream or downstream of the TSS of the target genes. For genes with multiple peaks, only one representative peak is shown. The distance to the TSS is shown.
